# NeutrobodyPlex - Nanobodies to monitor a SARS-CoV-2 neutralizing immune response

**DOI:** 10.1101/2020.09.22.308338

**Authors:** Teresa R. Wagner, Philipp D. Kaiser, Marius Gramlich, Elena Ostertag, Natalia Ruetalo, Daniel Junker, Julia Haering, Bjoern Traenkle, Matthias Becker, Alex Dulovic, Helen Schweizer, Stefan Nueske, Armin Scholz, Anne Zeck, Katja Schenke-Layland, Annika Nelde, Monika Strengert, Juliane S. Walz, Georg Zocher, Thilo Stehle, Michael Schindler, Nicole Schneiderhan-Marra, Ulrich Rothbauer

**Affiliations:** Pharmaceutical Biotechnology, Eberhard Karls University Tuebingen, Germany; Natural and Medical Sciences Institute at the University of Tuebingen, Germany; Interfaculty Institute of Biochemistry, Eberhard-Karls University Tuebingen, Germany; Institute for Medical Virology and Epidemiology of Viral Diseases, University Hospital, Tuebingen, Germany; Livestock Center of the Faculty of Veterinary Medicine, Ludwig Maximilians University Munich, Oberschleissheim, Germany; Cluster of Excellence iFIT (EXC2180) “Image-Guided and Functionally Instructed Tumor Therapies”, University of Tuebingen, Tuebingen, Germany; Department of Women’s Health, Research Institute for Womens’s Health, Eberhard-Karls-University, Tuebingen, Germany; Department of Medicine/Cardiology, Cardiovascular Research Laboratories, David Geffen School of Medicine at UCLA, Los Angeles, CA, USA; Clinical Collaboration Unit Translational Immunology, German Cancer Consortium (DKTK), Department of Internal Medicine, University Hospital Tuebingen, Tuebingen, Germany; Institute for Cell Biology, Department of Immunology, University of Tuebingen, Tuebingen, Germany; Dr. Margarete Fischer-Bosch Institute of Clinical Pharmacology and Robert Bosch Center for Tumor Disease, RBCT, Stuttgart, Germany; Department of Epidemiology, Helmholtz Centre for Infection Research, Braunschweig, Germany; TWINCORE GmbH, Centre for Experimental and Clinical Infection Research, a joint venture of the Hannover Medical School and the Helmholtz Centre for Infection Research, Hannover, Germany; Vanderbilt University School of Medicine, Nashville, TN, USA

**Author notes:** corresponding author: Prof. Dr. Ulrich Rothbauer, Natural and Medical Sciences Institute at the University of Tuebingen, Markwiesenstr. 55, 72770 Reutlingen, Germany.

## Abstract

As the COVID-19 pandemic escalates, the need for effective vaccination programs, therapeutic intervention, and diagnostic tools increases. Here, we identified 11 unique nanobodies (Nbs) specific for the SARS-CoV-2 spike receptor-binding domain (RBD) of which 8 Nbs potently inhibit the interaction of RBD with angiotensin-converting enzyme 2 (ACE2) as the major viral docking site. Following a detailed epitope determination and characterization of the binding mode by structural analysis, we constructed a hetero-bivalent Nb targeting two different epitopes within the RBD:ACE2 interface. This resulted in a high-affinity binder with a viral neutralization efficacy in the picomolar range. Using the bivalent Nb as a surrogate, we established a competitive multiplex binding assay (“NeutrobodyPlex”) for detailed analysis of the presence and performance of neutralizing RBD-binding antibodies in the serum of convalescent or vaccinated patients. As demonstrated, the NeutrobodyPlex enables high-throughput screening and detailed analysis of neutralizing immune responses in infected or vaccinated individuals, helping to monitor immune status or guide vaccine design. This approach is easily transferrable to diagnostic laboratories worldwide.

## Introduction

By the end of 2020, the COVID-19 pandemic has caused the death of more than 1.6 million people worldwide and the current lack of a cure or established vaccine comes with severe lockdowns and dramatic economic losses. Neutralizing antibodies targeting the causative agent of the disease, severe acute respiratory syndrome coronavirus 2 (SARS-CoV-2), are of substantial interest for prophylactic and therapeutic options and help to guide vaccine design^1^. These antibodies bind to the receptor-binding domain (RBD) of the homotrimeric transmembrane spike (spike) and prevent viral entry via the angiotensin-converting enzyme II (ACE2) expressed on human epithelial cells of the respiratory tract^2^. Since the outbreak of the pandemic, a constantly growing number of neutralizing antibodies has been identified from COVID-19 patients, underlining the importance of RBD-specific antibodies in blocking the RBD:ACE2 interaction site for the development of a protective immune response^3-8^. A promising alternative to conventional antibodies (IgGs) are single-domain antibodies (nanobodies, Nbs) derived from the heavy-chain antibodies of camelids (**Figure 1**)^9^. Due to their small size and compact fold, Nbs show high chemical stability, solubility and fast tissue penetration. Nbs can be efficiently selected against different epitopes on the same antigen and can be easily converted into multivalent formats^9^. The potential of Nbs to address SARS-CoV-2 has been impressively demonstrated by the recent identification of several RBD specific Nbs from naïve/ synthetic libraries^10-12^ or immunized animals^12-16^.

**Figure 1:**
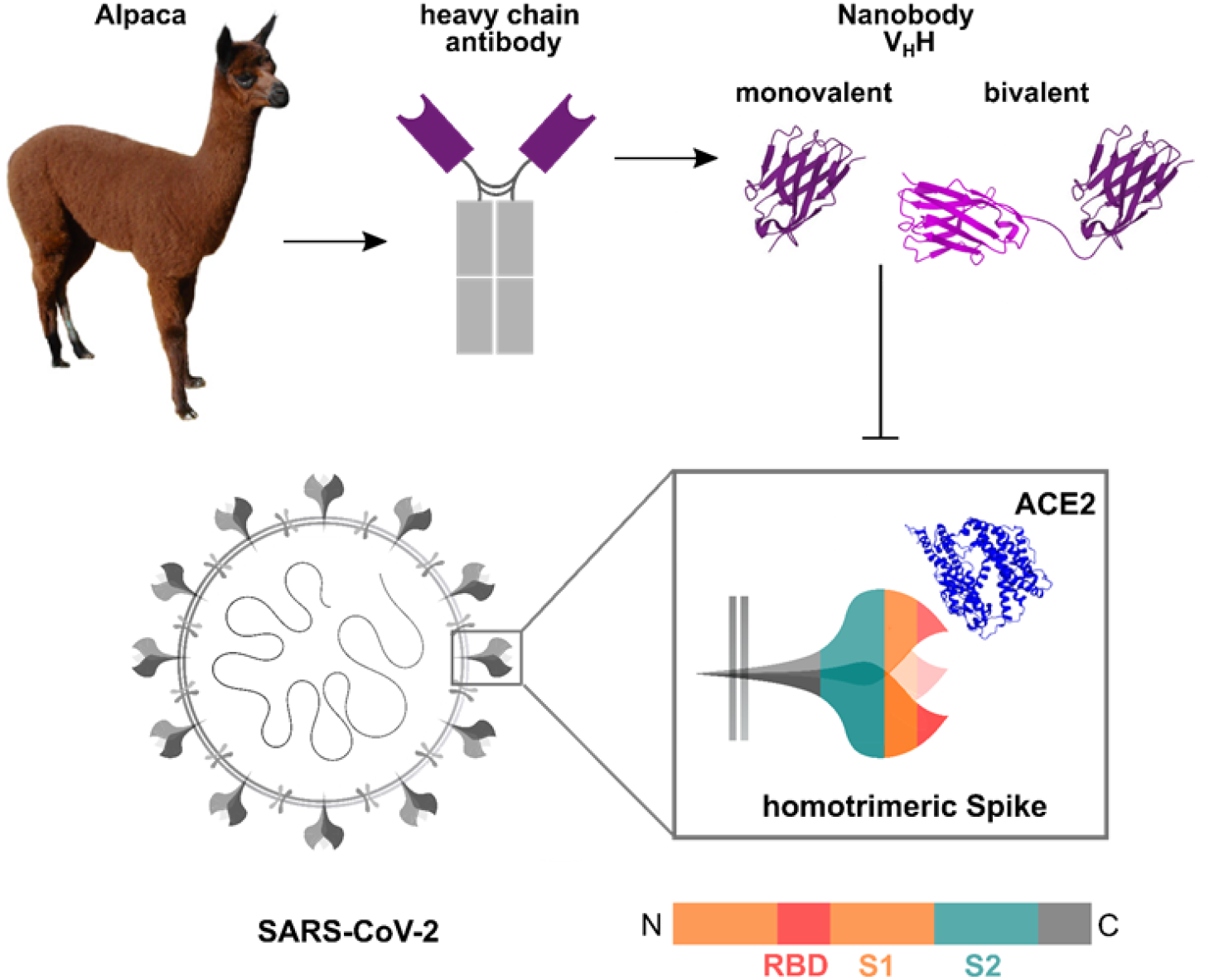
Generation of nanobodies blocking the SARS-CoV-2 RBD:ACE2 interface. Nanobodies (Nbs) are genetically engineered from heavy-chain only antibodies of alpacas. The interaction between the SARS-CoV-2 homotrimeric spike protein and angiotensin converting enzyme (ACE) 2 can be blocked by receptor-binding domain (RBD)-specific Nbs in the monovalent or bivalent format. Protein structures adapted from PDB 3OGO (Nb)^20^ and 6CS2 (ACE2)^21^.

Since the pandemic outbreak multiple serological SARS-CoV-2 assays have been established to monitor seroconversion in individuals and estimate the level of endemic infection in the general population. However, most available serological tests measure the full immune response and can therefore not differentiate between total binding and neutralizing antibodies^4,17,18^. Detection of the latter is still mostly performed by conventional virus neutralization tests (VNTs), which are both time consuming (2 - 4 days) and require work with infectious SARS-CoV-2 virions in a specialized biosafety level 3 (BSL3) facility^19^. To overcome these limitations, we aimed to employ Nbs as antibody surrogates and developed a competitive binding approach to screen for neutralizing antibodies on a high-throughput basis in samples from patients or vaccinated individuals.

Here we describe the selection of 11 unique Nbs derived from an alpaca immunized with glycosylated SARS-CoV-2 RBD. Employing a multiplex *in vitro* binding assay, detailed epitope mapping and structural analysis, we identified 8 Nbs that effectively block the interaction between RBD, S1-domain (S1) and the homotrimeric spike protein (spike) with ACE2 and neutralize SARS-CoV-2 infection in a human cell line. Generation of a hetero-bivalent Nb (bivNb) simultaneously targeting different epitopes within the RBD:ACE2 interface results in a potent antibody surrogate with substantially improved binding affinities and functional IC50 values in the low picomolar range. To monitor the presence and performance of neutralizing antibodies addressing the RBD:ACE2 interface in convalescent patient samples, we implemented the bivNb in a competitive multiplex binding assay, termed NeutrobodyPlex. Based on the data presented in this study, the NeutrobodyPlex provides a versatile high-throughput approach to screen for a neutralizing immune response in infected or vaccinated individuals, helping to monitor immune status of large populations, outcome of vaccination campaigns and to guide vaccine design.

## Results

### Selection of SARS-CoV-2-specific Nbs

To generate Nbs directed against the RBD of SARS-CoV-2 we immunized an alpaca (*Vicugna pacos*) with purified RBD^22^ and established a Nb phagemid library comprising ∼4 x 10^7^ clones representing the full repertoire of variable heavy chains of heavy-chain antibodies (V_H_Hs or Nbs). After two rounds of phage display on passively adsorbed or biotinylated RBD immobilized on streptavidin plates, we analyzed 492 individual clones in a solid-phase phage ELISA and identified 325 positive binders. Sequence analysis of 72 clones revealed 11 unique Nbs which cluster into eight families displaying highly diverse complementarity determining regions (CDR) 3 (**Figure 2 a**). Individual Nbs were produced and purified from *Escherichia coli (E*.*coli)* (**Figure 2 b**) and affinity measurements revealed K_D_ values ranging from ∼1.3 nM to ∼53 nM indicating the selection of 10 high affinity monovalent binders, whereas NM1225 displaying a binding affinity in the micromolar range was not considered for further analysis (**Figure 2 c, Supplementary Figure 1**). Next, we analyzed whether the selected Nbs can block the interaction between the isolated receptor-binding domain (RBD), S1 domain (S1) or homotrimeric spike protein (spike) of SARS-CoV-2 to the human ACE2 receptor. To this end, we utilized a multiplex ACE2 competition assay employing the respective SARS-CoV-2 antigens coupled to paramagnetic beads (MagPlex Microspheres)^23^ and incubated them with biotinylated ACE2 and dilutions of purified Nbs before measuring the residual binding of ACE2. Additionally, we included a non-specific Nb (GFP-Nb) as negative control and two inhibiting mouse antibodies^24^ as positive controls. Data obtained through this multiplex binding assay showed that 8 of the 10 analyzed Nbs inhibit ACE2 binding to all tested SARS-CoV-2 antigens (**Supplementary Figure 2**). Notably, IC_50_ values obtained for the most potent inhibitory Nbs NM1228 (0.5 nM), NM1226 (0.85 nM) and NM1230 (2.12 nM) are highly comparable to IC_50_ values measured for the mouse IgGs (MM43: 0.38 nM; MM57: 3.22 nM).

**Figure 2:**
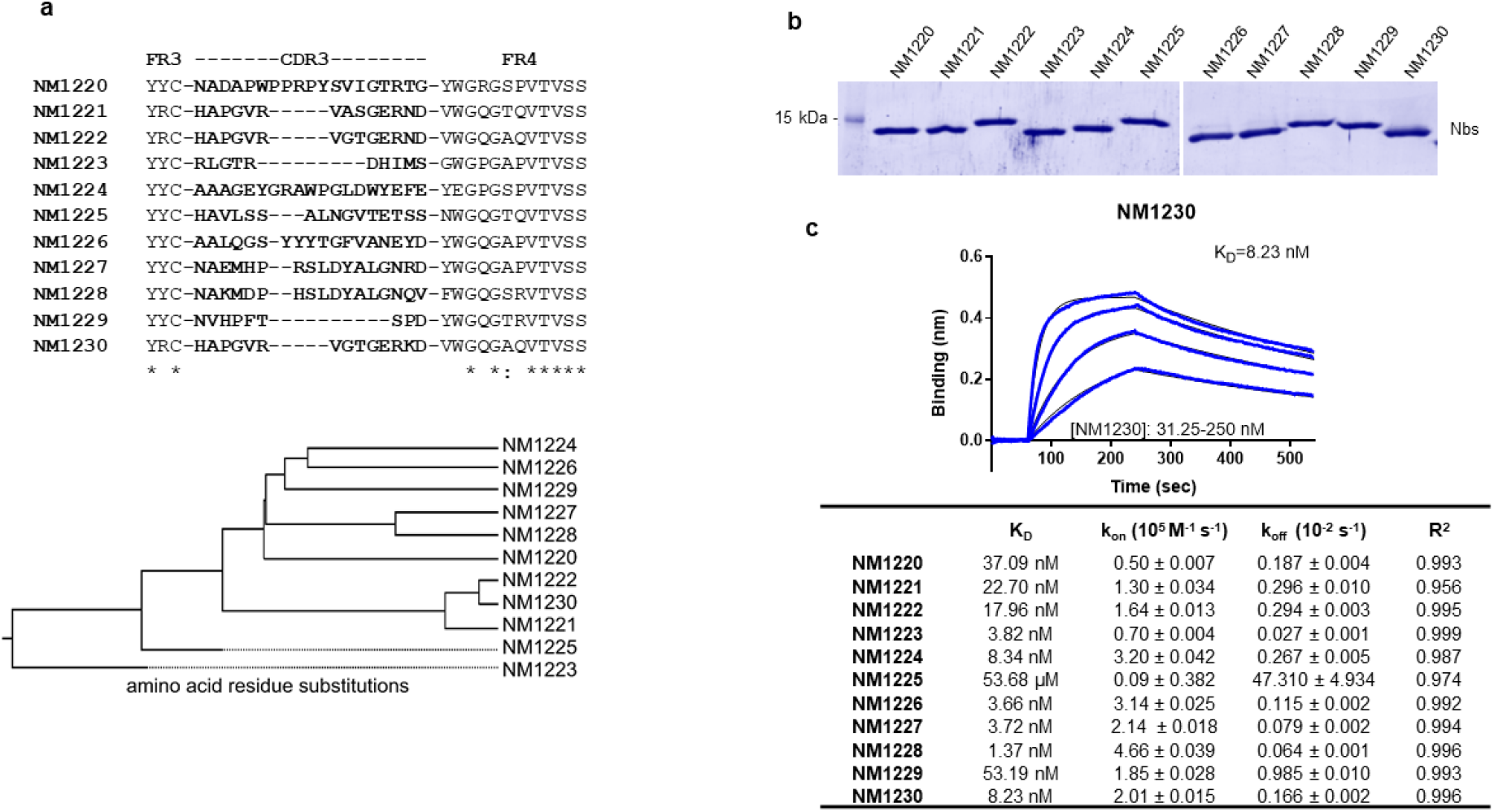
Biochemical characterization of RBD-specific Nbs. (**a**) Amino acid sequences of the complementarity determining region (CDR) 3 from unique Nbs selected after two rounds of biopanning are listed (upper panel). Phylogenetic tree based on a ClustalW alignment of the Nb sequences is shown (lower panel). (**b**) Recombinant expression and purification of Nbs using immobilized metal affinity chromatography (IMAC) and size exclusion chromatography (SEC). Coomassie staining of 2 µg of purified Nbs is shown. (**c**) For biolayer interferometry based affinity measurements, biotinylated RBD was immobilized on streptavidin biosensors. Kinetic measurements were performed by using four concentrations of purified Nbs ranging from 15.6 nM - 2 µM. As an example, the sensogram of NM1230 at indicated concentrations is shown (upper panel). The table summarizes affinities (K_D_), association (K_on_) and dissociation constants (K_off_) determined for individual Nbs (lower panel).

### Epitope Identification

To identify the relative location of Nb epitopes Nbs within the RBD, we firstly performed epitope binning experiments of Nb combinations using biolayer interferometry. For this, sensors, pre-coated with biotinylated RBD, were loaded with a Nb from one family followed by a short dissociation step and subsequent loading of a second Nb from a different family (**Supplementary Figure 3 a**). As expected, Nbs displaying similar CDR3 sequences (NM1221, NM1222 and NM1230, Nb-Set2) were unable to bind simultaneously as they recognize identical or highly similar epitopes. Interestingly, we noticed that Nbs with highly diverse CDR3s such as NM1228, NM1226, NM1227 and NM1229 also could not bind simultaneously, suggesting that these Nbs also recognize similar or overlapping epitopes. Consequently, we clustered these diverse Nbs into Nb-Set1. In total, we identified five distinct Nbs-Sets, comprising at least one candidate targeting a different epitope within the RBD compared to any member of a different Nb-Set (**Supplementary Figure 3 b**). For detailed epitope mapping, we performed Hydrogen-Deuterium Exchange Mass Spectrometry (HDX-MS) using the most potent inhibitory Nbs selected from the different Nb-Sets. Both members of Nb-Set1, NM1226 and NM1228, interacted with the RBD at the back/ lower right site (Back View, **Figure 3**). The highest exchange protection for NM1226 was found in the amino acid (aa) region N370 – L387 which is also covered by NM1228. Additionally, NM1228 binds to aa Y489 - S514, being part of the RBD:ACE2 binding interface. NM1230 (Nb-Set2) shows the highest exchange protection ranging from C432 - L452 covering two amino acids involved in ACE2 binding (G446, Y449). Additionally, a second protected region was found covering aa N487 - G496 with overlap to the RBD:ACE2 interface (**Figure 3**). In accordance to our binning studies, the main epitope regions differ from both Nb-Sets. As expected, NM1221 and NM1222 (both Nb-Set2) addressed similar RBD regions compared to NM1230 (**Supplementary Figure 4**) while NM1224 (Nb-Set4) showed an interaction distinct from all other Nbs, covering both its main binding region located at the lower right side (**Supplementary Figure 4**, Front View) and residues in the ACE2:RBD interface (**Figure 3**, Front View, upper left corner). Additionally, the non-inhibitory NM1223 (Nb-Set3) (**Supplementary Figure 4**) was shown not to contact any amino acid residues involved in the RBD:ACE2 interface but rather binds to the opposite side (**Supplementary Figure 4**, Front View).

**Figure 3:**
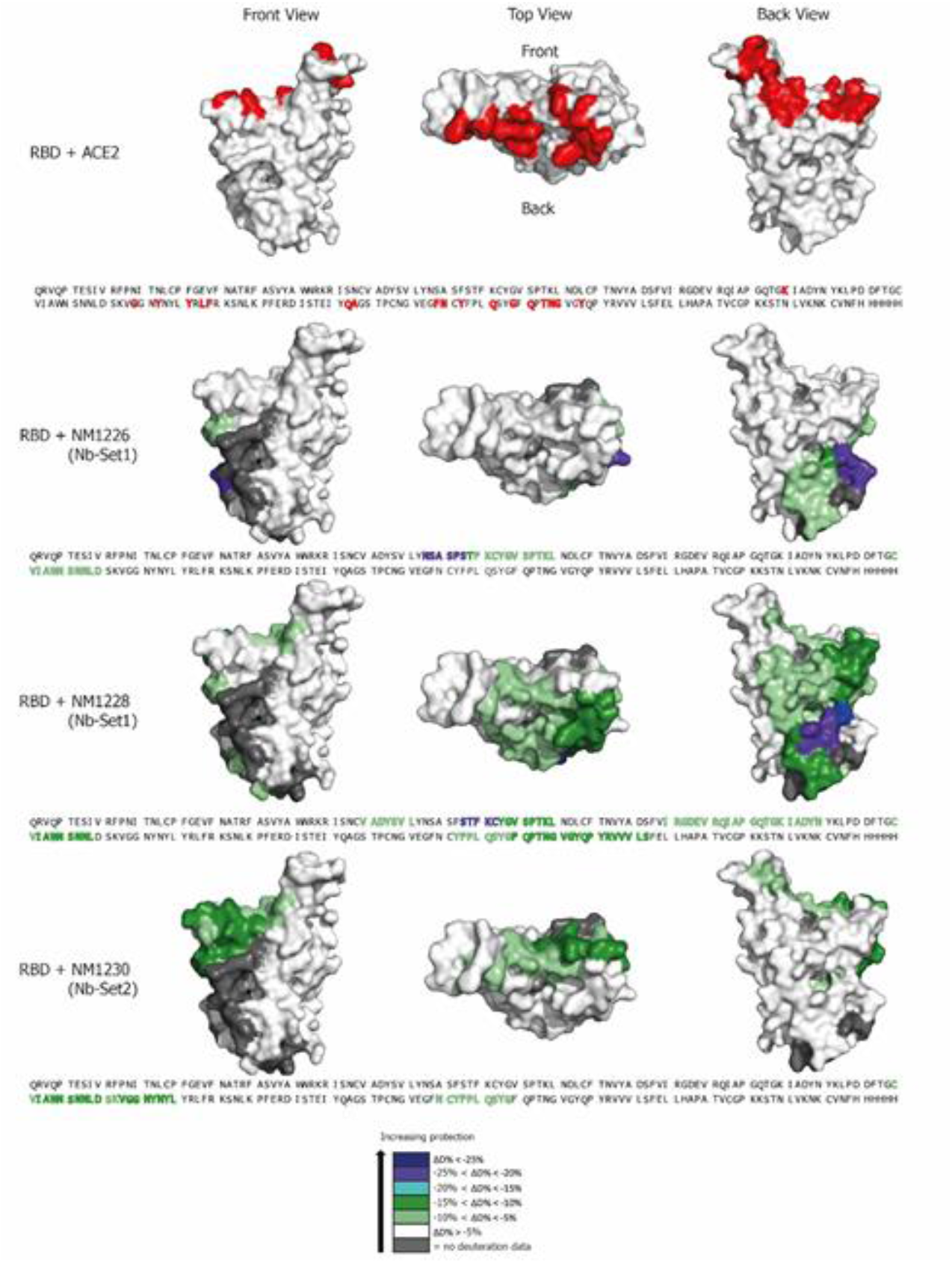
Epitope mapping of Nbs by HDX mass spectrometry. Surface structure model of RBD showing the ACE2 interface and the HDX-MS epitope mapping results of NM1226, NM1228 and NM1230. Amino acid residues of RBD (PDB 6M17 ^2^) involved in the RBD:ACE2 interaction site^2,25^ are shown in red (top panel). RBD epitopes protected upon Nb binding are highlighted in different colors indicating the strength of protection. Accordingly amino acid residues which are part of the Nb recognized epitopes are highlighted in the RBD sequence

### Crystal structure of RBD:NM1230 complex

To obtain a deeper insight into the binding mode of one of the most potent inhibitory Nbs, we solved the crystal structure of NM1230 in complex with RBD at a resolution of 2.9 Å (**Supplementary Table 1**). Analysis of the RBD:NM1230 complex revealed that the main contact is formed by residues located in the CDR3 of NM1230 (residues P99, R102 E107, R108, K109, D110, V111, W112). Additionally, the N-terminus (Q1), framework region 2 (FR2, residues N35, Y37, Q39, G42, K43, A44, A49, L45 and L47) and FR3 (residue E61) of NM1230 form contact sites. On the RBD site residues that interact with NM1230 are Y351 and the loop region formed by residues 437 - 503 including Y453, Y449, N450, L468, T470, E484, Y489, F490, L492, N493, and S494) (**Figure 4 a**). The RBD:NM1230 interface RBD is mainly formed by polar contacts and one salt-bridge (R_NM1230_102 to E_RBD_484), but π-π stackings (W112_NM1230_ and F490_RBD_, R108_NM1230_ to Y489_RBD_) are also observed (**Figure 4 b**). In total it buries a surface area of ∼830 Å^2^, which is comparable (∼860 Å^2^) to the contact interface in recently reported structure of the SARS-CoV-2 spike protein bound to neutralizing Nb-Ty1^13^. Superposition of the RBD:NM1230 complex structure with the spike protein in the complex with Nb-Ty1 reports a C_α_-rmsd deviation of 1.2 Å for all three RBD domains showing that the binding sites of Nbs NM1230 and Nb-Ty1 partially overlap (**Figure 4 c**). Nevertheless, subtle changes establish an alternative binding epitope for NM1230 (**Figure 4 c, Supplementary Figure 5 a**). Next, we compared the RBD:NM1230 structure with the recently reported RBD:ACE2 receptor complex^2^ to structurally validate the neutralizing potential of NM1230. Closer inspection of the binding site (**Supplementary Figure 5 b**) showed that NM1230 only partially overlaps with the ACE2 binding interface and that the neutralizing effect can be isolated to a limited set of residues within the RBD (K417, Y449, F456, Y489, Q493, S494) (**Supplementary Figure 5**). However, NM1230 does not only block binding to ACE2 via its interaction to the RBD on the same protomer (**Figure 4 c, clash I**), but also impairs ACE2 binding through steric collision on the neighbouring RBD (**Figure 4 c, clash II**). Based upon our structural data, we propose that NM1230-mediated blocking of two out of three RBDs would suffice to abolish ACE2 binding to a trimeric spike molecule.

**Figure 4:**
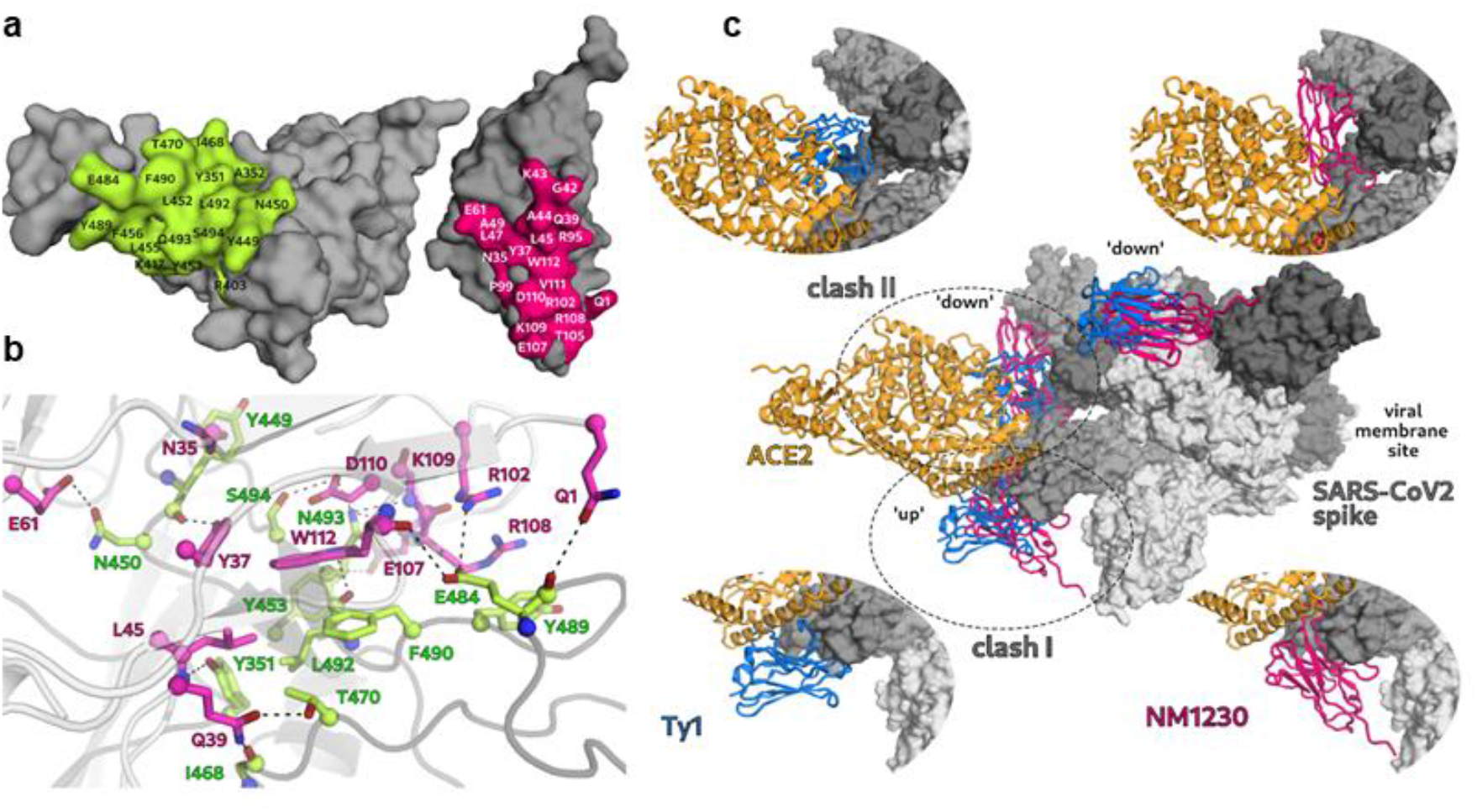
Structure of RBD:NM1230 complex. (a) The binding interface of RBD (green) and NM1230 (pink) lists all residues involved in contact formation. (b) Close-up view of the interface, with direct interactions of NM1230 (pink) and RBD (green). (c) Alignment of the SARS-CoV-2 spike:Ty1-Nb (blue) cryoEM structure13 with the RBD:NM1230 complex. The ACE2 (orange) and the NM1230 (pink) and Ty1-Nb (blue) are depicted as cartoons, whereas the spike trimer is shown in surface representation (grey). The neutralizing effect of NM1230 and Ty1-Nb is shown by expected collisions (clash I and clash II) with ACE2 in the alignment. Additional close up views of the individual Nb positions are shown to highlight the differences in Nb binding and ACE2 blocking.

### Nanobodies show a high potency in neutralizing SARS-CoV-2

For functional analyses, we screened the Nbs for their virus-neutralizing potencies. For a viral neutralization test (VNT), human Caco-2 cells were co-incubated with the SARS-CoV-2-mNG infectious clone and serial dilutions of NM1223, NM1224, NM1226, NM1228, NM1230 or GFP-Nb as negative control. 48 h post-infection neutralization potency was determined via automated fluorescence-microscopy of fixed and nuclear-stained cells (**Supplementary Figure 6**). The infection rate normalized to a non-treated control was plotted and IC_50_ values were determined via sigmoidal inhibition curve fits (**Supplementary Figure 6**). NM1226 and NM1228 (both Nb-Set1), showed the highest neutralization potency with IC_50_ values of ∼15 nM and ∼7 nM followed by NM1230 (Nb-Set2; ∼37 nM) and NM1224 (Nb-Set4; ∼256 nM). As expected, NM1223 (Nb-Set3) was not found to neutralize SARS-CoV-2 (**Supplementary Figure 6**). Overall, these findings are highly consistent compared to our results obtained from the multiplex ACE2 competition assay, thus demonstrating high potencies of ACE2-blocking Nbs to neutralize viral infection.

### Generation of a hetero-bivalent Nb with improved efficacies

Having identified a range of Nbs with high potency, inhibiting and neutralizing characteristics, we proposed that the most potent of these candidates derived from Nb-Set1 (NM1226) and Nb-Set2 (NM1230) might act synergistically. To examine this, we genetically fused the coding sequences head-to-tail by a flexible Gly-Ser linker and generated a hetero-bivalent Nb (bivNb) named NM1267 (NM1230/NM1226). Following production and purification from mammalian cells (**Figure 5 a**), we determined the affinity (**Figure 5 b, c**) and analyzed its performance in our multiplex ACE2 competition assay. The bivNb showed a considerably improved affinity and revealed an outstanding inhibition of ACE2 binding to RBD, S1 and spike with an IC_50_ in the low picomolar range (**Figure 5 b, d)**. Additionally, we observed an increased potency for viral neutralization indicated by an IC_50_ of ∼0.9 nM (**Figure 5 b, e**), demonstrating that the bivalent NM1267 simultaneously targeting two different epitopes within the RBD:ACE2 interaction site, embodies a substantially refined tool which is highly beneficial for viral neutralization and competitive binding studies.

**Figure 5:**
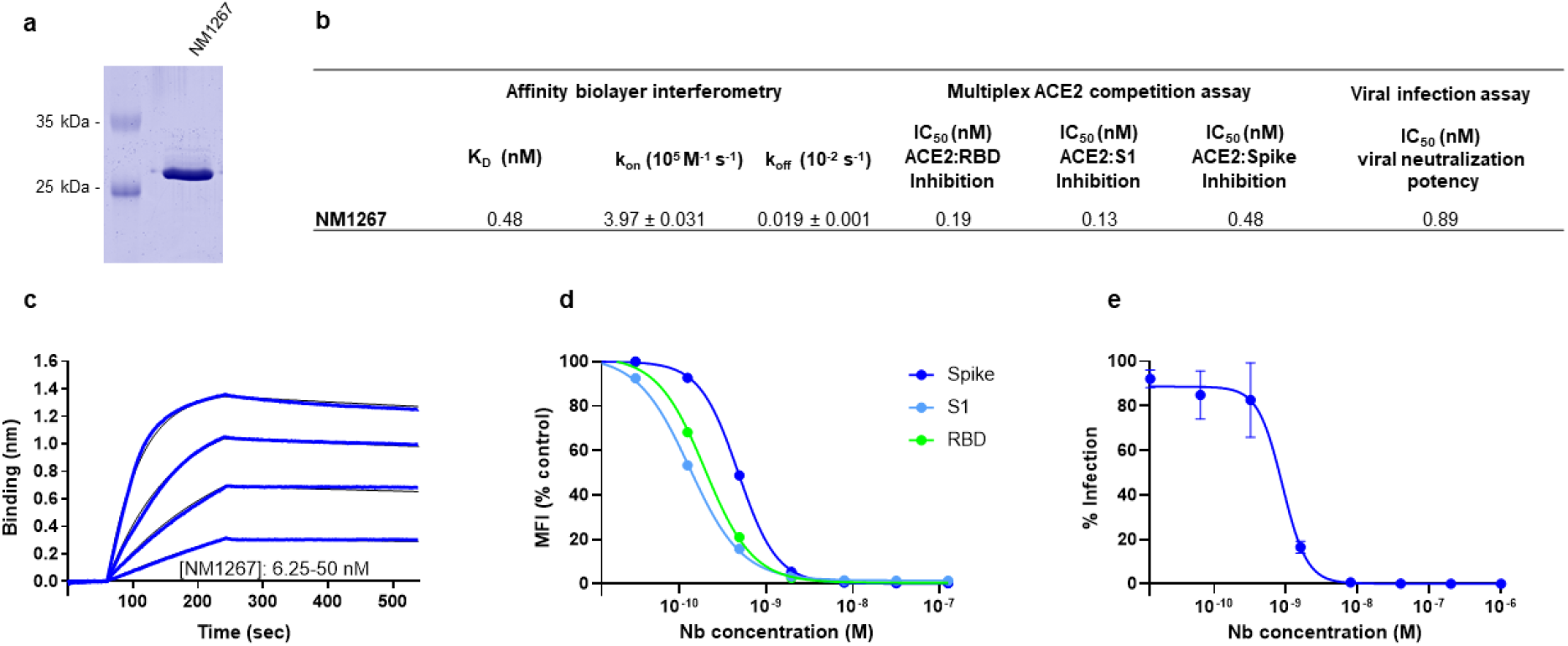
Hetero-bivalent NM1267 competes with ACE2 and neutralizes SARS-CoV-2 infection. (**a**) Coomassie staining of 1 µg of purified hetero-bivalent Nb NM1267. (**b**) Table summarizing the affinity (K_D_) determined by biolayer interferometry and IC_50_ values of the multiplex ACE2 competition assay and viral infection assay obtained for the bivNb NM1267. (**c**) Sensogram of affinity measurements via biolayer interferometry using four concentrations (6.25 - 50nM) of NM1267. (**d**) Results from multiplex ACE2 competition assay are shown for the three spike-derived antigens: RBD, S1-domain (S1) and homotrimeric spike (Spike). NM1267 was diluted from 126 nM to 7.69 pM in the presence of 80 ng/mL ACE2 and residual antigen-bound ACE2 was measured. MFI signals were normalized to the maximum detectable signal per antigen given by the ACE2-only control. IC_50_ values were calculated from a four-parametric sigmoidal model. Data are presented as mean +/- SD of three technical replicates (n = 3). (**e**) Neutralization potency of NM1267 was analyzed in Caco-2 cells using the SARS-CoV-2-mNG infectious clones. Infection rate normalized to virus-only infection control is illustrated as percent of infection (% Infection). IC_50_ value was calculated from a four-parametric sigmoidal model and data are presented as mean +/- SD of three biological replicates (N = 3).

### NeutrobodyPlex – Using Nbs to determine a SARS-CoV-2 neutralizing immune response

Currently available serological assays provide data on the presence and distribution of antibody subtypes against different SARS-CoV-2 antigens within serum samples of infected and recovered SARS-CoV-2 patients^4,17,22,23,26^. However, they do not differentiate between total and neutralizing RBD binding antibodies, which sterically inhibit viral entry via ACE2^7,8^. To address this important issue, our bivNb NM1267 might be a suitable surrogate to monitor the emergence and presence of neutralizing antibodies in serum samples of patients. We speculated that NM1267 specifically displaces such antibodies from the RBD:ACE2 interface, which can be monitored as a declining IgG signal (**Figure 6 a**). To test this hypothesis, we generated a high-throughput competitive binding assay, termed NeutrobodyPlex, by implementing the NM1267 in a recently developed, multiplexed serological assay^23^. We co-incubated antigen-coated beads comprising RBD, S1 or spike with serum samples from 5 patients and a dilution series of NM1267. Mean fluorescent intensities (MFI) derived from antigen-bound IgGs in the presence of bivNb were normalized to the MFI values of IgGs in the serum-only samples, illustrated as MFI (% control). When analyzing IgG binding to RBD, we detected a complete displacement in the presence of ∼63 nM NM1267. Similarly, a distinct signal reduction for S1 binding IgGs became observable reaching a plateau upon addition of ∼63 nM NM1267 (**Figure 6 b, Supplementary Table 2**). Notably, we observed only minor signal reduction when analyzing spike binding IgGs (**Figure 6 b**) indicating that the majority of serum IgGs bind this large antigen at epitopes beyond the RBD:ACE2 interaction site. From this data, we concluded, that all 5 tested individuals comprise a substantial fraction of neutralizing IgGs, which can be detected by competitive bivNb binding using RBD or S1 as antigens. To demonstrate that the NeutrobodyPlex further determines the presence of neutralizing IgGs in detailed resolution, we highlight the results from monitoring the displacement of S1-binding IgGs in two selected patient samples #289 and #265. Upon addition of increasing concentrations of NM1267, we observed a prominent signal reduction of ∼ 85% in patient #289 whereas samples of patient #265 only revealed ∼ 53% displacement of S1-binding IgGs (**Figure 6 c**).

**Figure 6:**
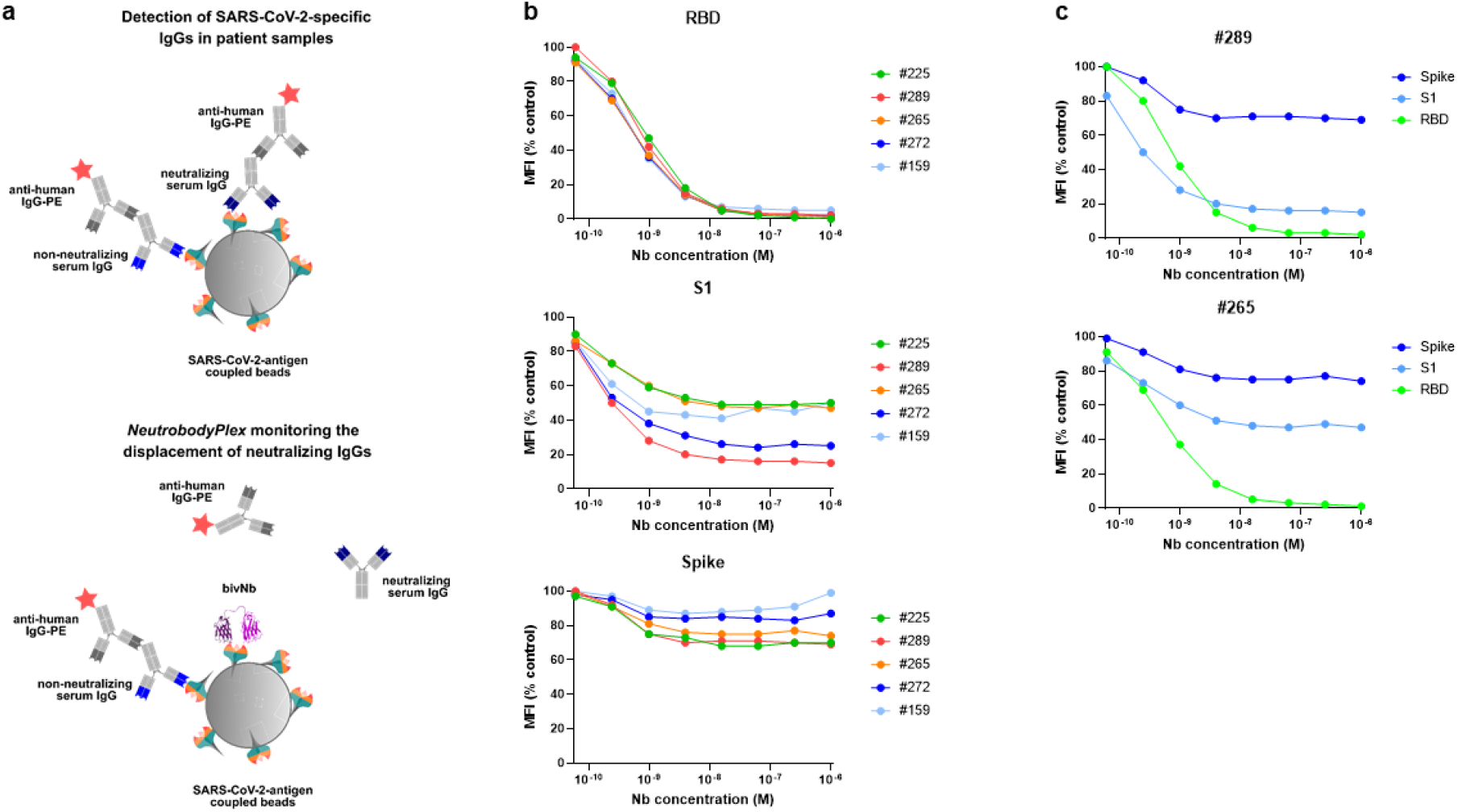
NeutrobodyPlex - multiplex competitive binding assay to monitor a neutralizing immune response in patients. (**a**) Schematic illustration of the NeutrobodyPlex. The displacement of serum-derived neutralizing IgGs binding to SARS-CoV-2 antigens upon addition of bivNb is measured. In presence of neutralizing IgGs, the fluorescent signal from anti-human-IgG-PE is inversely proportional to the applied bivNb concentration. (**b**) For the NeutrobodyPlex, antigen-coated beads comprising RBD, S1 or spike were co-incubated with serum samples from 5 patients and a dilution series of NM1267 (1 µM to 6 pM). Mean fluorescent intensities (MFI) derived from antigen-bound IgGs in the presence of bivNb normalized to the MFI values of IgGs in the serum-only samples, illustrated as MFI (% control), are shown. (**c**) For two patient derived serum samples (#289, #265), differences in the NM1267-mediated displacement of IgGs binding the three spike-derived antigens (RBD, S1, spike) are shown.

Next, we compared the NeutrobodyPlex on RBD with the cell-based VNT by analyzing a set of 18 serum samples from convalescent SARS-CoV-2 patients, collected on days 19 - 57 following a positive PCR test result, and 4 control samples from healthy donors. To detect differences within the overall immune response and to qualify the neutralizing capacity of the serum samples, we performed the NeutrobodyPlex using two concentrations of NM1267 previously shown to completely (1 µM) or partly (1 nM) displace IgGs binding the RBD:ACE2 interface (**Figure 6 b**). Both assays (**Figure 7**) revealed the presence of neutralizing antibodies in all serum samples from convalescent individuals, as seen when using the highest (1:40) serum dilution in the viral infection assay or upon the addition of 1 µM NM1267 in the NeutrobodyPlex. Notably, neutralizing IgGs were not detected in any of the control serum samples (**Supplementary Table 3**). When further analyzing the displacement of RBD binding IgGs using the lower NM1267 (1 nM) concentration, substantial variations in individual neutralizing capacities became visible. While some patient samples comprised potent, high affinity neutralizing IgGs which were able to outcompete NM1267 for binding to the RBD:ACE2 interface (**Figure 7 a**, high MFI (% control), light-colored squares), a continuous bivNb-mediated displacement of IgGs was detectable in other samples (**Figure 7 a**, low MFI (% control) dark-colored squares) indicating the presence of IgGs with a lower neutralizing potency. In parallel, the VNT using higher serum dilutions also revealed substantial differences in the individual neutralizing potencies (**Figure 7 b**). To confirm the validity of the NeutrobodyPlex, we calculated the mean percent of infection (% Infection) of all individual serum dilutions and plotted these values against the respective MFI (% control) obtained from the NeutrobodyPlex (**Figure 7 c**). The observed negative correlation (high MFI (% control) vs low mean % Infection) strongly suggests that the NeutrobodyPlex allows the detection of individual neutralizing immune responses and assessment of their potency.

**Figure 7:**
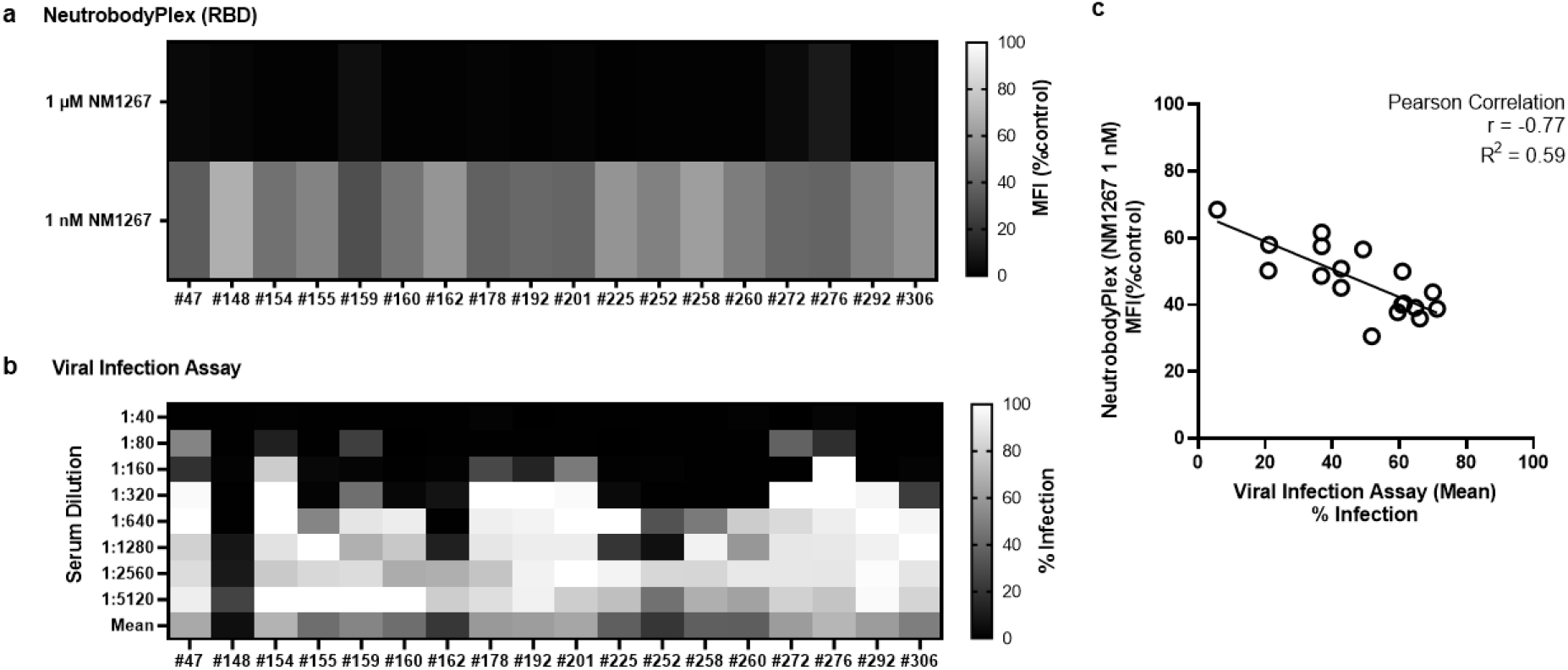
Specificity of the NeutrobodyPlex in determining individual neutralizing immune responses. Serum samples of 18 convalescent SARS-CoV-2-infected individuals were analyzed using the NeutrobodyPlex and the viral infection assay. (**a**) For the NeutrobodyPlex on RBD, two concentrations of NM1267 (1 µM and 1 nM) were applied. Light-colored squares (high MFI(%control)) are indicative for IgGs outcompeting NM1267 from the RBD:ACE2 interface, dark-colored squares (low MFI (%control) show a continuous displacement of IgGs from serum samples in the presence of NM1267. (**b**) For the viral infection assay, serial dilutions of the serum samples (1:40 - 1:5120) were applied. Dark colored squares are indicative for a low infection level (low % Infection), light colored squares show a high infection rate (high % Infection). (**c**) The mean percent of infection (% Infection) derived from all individual serum dilutions obtained by the viral infection assay was calculated and plotted against the respective MFI (% control) obtained from the NeutrobodyPlex on RBD in the presence of NM1267 (1 nM). The Pearson correlation determined a negative correlation with r = -0.77 and R^2^ = 0.59 (p <0.05).

For final validation, we screened a serum sample set of 112 convalescent SARS-CoV-2-infected and 8 uninfected individuals in the NeutrobodyPlex. In addition to RBD, S1 and spike, we included the S2 domain (S2) and the nucleocapsid (N) of SARS-CoV-2 as controls. Incubation with the higher amount of NM1267 (1 µM) revealed the presence of neutralizing IgGs in all serum samples from convalescent patients as shown by an effective replacement of IgGs from RBD, S1 and partially from spike, but not from S2 or N (**Supplementary Figure 7 a**). In line with the findings from the initial patient sample set, differences in the potencies of neutralizing IgGs present in the individual serum samples became detectable upon addition of the lower concentration of NM1267 (1 nM) (**Supplementary Figure 7 b**). As observed previously, non-infected individuals did not display any detectable antibody signal against the presented SARS-CoV-2 antigens and experienced no alteration upon NM1267 addition (data not shown). We further analyzed if the total level of SARS-CoV-2 specific IgGs correlates with the amount of neutralizing IgGs. As a marker for total IgGs we defined MFI values measured for spike-binding IgGs and plotted them against normalized MFI values from IgGs binding to RBD in the presence of both concentrations (1 µM, 1 nM) of NM1267 (MFI RBD (%control)) (**Figure 8**). We found high variability of total IgGs (MFI: ∼3500 – ∼47000) in the tested serum samples. Upon addition of the higher concentration of NM1267 (1 µM), we observed a complete displacement of IgGs binding the RBD:ACE2 interface independent of the amount of total IgG (**Figure 8**). Notably, no measurable correlation between the total IgG and the displaced IgG signal was detected in the presence of the lower amount of bivNbs (**Figure 8**). According to our data, there might be an increased likelihood for the presence of neutralizing antibodies in patients showing an overall high SARS-CoV-2 specific IgG level, however as illustrated in two examples (**Figure 8**, highlighted in green) this cannot be generally assumed. While sample #36 comprises only a minor amount of total IgGs (MFI spike (control) ∼3700) we measured a high neutralizing potency, as displayed by the displacement of only ∼25% of RBD binding IgGs in the presence of NM1267 (1 nM). In contrast, a high amount of total IgGs was detected in sample #250 (MFI spike (control) ∼32,000) but ∼70% of RBD-specific IgGs were continuously displaced by NM1267 (1 nM) (**Figure 8**) indicating the presence of low levels and/or low affine neutralizing IgGs in this individual. In summary, these findings show how the NeutrobodyPlex can provide detailed information not only on the presence of neutralizing antibodies in patient samples, but also enable a qualitative and quantitative assessment of the individual immune response.

**Figure 8:**
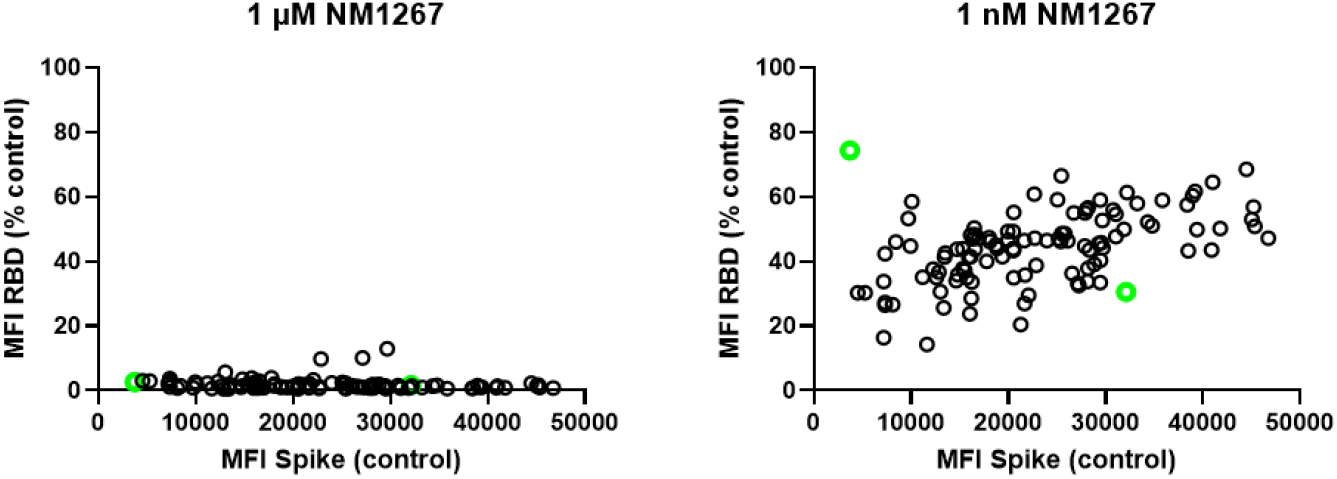
NeutrobodyPlex enables a differentiated analysis on neutralizing IgGs compared to total SARS-CoV-2 binding IgGs in individuals. To investigate neutralizing capacities in relation to total levels of SARS-CoV-2 specific IgGs in individuals, serum samples of 112 convalescent SARS-CoV-2-infected individuals were analyzed using the NeutrobodyPlex. As a marker for total IgGs we defined MFI values measured for spike-binding IgGs (MFI Spike (control)) and plotted them against normalized MFI values from IgGs binding to RBD (MFI RBD (%control)) in the presence of both concentrations of NM1267 (1 µM, left panel; 1 nM, right panel). Two patient samples (#36 and #250) displaying strikingly different neutralizing potencies are highlighted as green circles.

## Discussion

Indisputably, advanced diagnostic tools and therapeutics against SARS-CoV-2 infection are desperately needed. Several studies have demonstrated that humoral immune responses giving rise to antibodies targeting the interaction site of RBD and the ACE2 receptor exposed on human epithelial cells of the respiratory tract result in a strong viral neutralizing effect^7,8,27,28^. Consequently, there is a strong demand to expand currently applied acute serological diagnostics measuring the overall immune response against SARS-CoV-2, towards a more differentiated analysis to specifically monitor and classify the emergence and presence of neutralizing antibodies in tested individuals. To monitor the presence of neutralizing antibodies addressing the RBD:ACE2 interface in serum samples, we developed a set of RBD-specific Nbs and screened for effective surrogates of RBD-binding IgGs. Of these, we selected high-affinity binding candidates which blocked the interaction between RBD and ACE2 in the context of different SARS-CoV-2 spike-derived antigens and additionally displayed a high viral neutralizing potency. Following detailed epitope mapping and structural analysis, we identified two binders which target different epitopes within the RBD:ACE2 interface and converted them into a hetero-bivalent Nb (NM1267). The bivalent NM1267 showed excellent performance in our multiplex ACE2 competition assay and the viral infection test with IC_50_ values in picomolar range. Notably, data from our biochemical analyses and the viral neutralization test are in good accordance, which strongly suggests that the multiplex ACE2 competition assay as presented is highly relevant and suitable to identify virus-neutralizing binders. For detecting a neutralizing immune response in serum samples of SARS-CoV-2 infected individuals, we implemented the bivalent NM1267 as a most potent neutralizing IgG surrogate in our previously described multiplex immunoassay (MULTICOV-AB^23^) and developed a novel diagnostic test called NeutrobodyPlex. Using NM1267 to specifically displace IgGs from binding to the RBD:ACE2 interface, we provide an in-depth analysis of the neutralizing immune response in a large patient cohort. The NeutrobodyPlex was successful in not only detecting the overall presence of neutralizing IgGs targeting the RBD:ACE2 interface, but could also categorize them according to their potencies. Additionally, the NeutrobodyPlex enables the comparison between neutralizing IgGs and the total level of SARS-CoV-2 antibodies, which will be an important parameter for determining the success of a vaccine.

In comparison to recently described assays directly measuring antibody-derived displacement of ACE2 in order to determine a neutralizing immune response^29^ a distinct lower binding affinity of ∼15 nM of RBD to ACE2 has to be considered. As a result, a neutralizing immune response may be detected even in the presence of weak RBD-binding antibodies, which could lead to a false statement of a protective immune response. The use of small-sized antibody surrogates exclusively binding to the RBD:ACE2 interface lowers the possibility of a non-targeted and non-reproducible displacement of ACE2 e.g. mediated by Fc regions of non-specifically binding IgGs. Finally, as demonstrated, the NeutrobodyPlex is performed in a bead-based multiplex setting, thus allowing to easily expand the antigen panel to accommodate e.g. novel, relevant RBD mutants occurring during the pandemic and which are highly relevant to be included in advanced vaccination strategies.

To our knowledge, the NeutrobodyPlex employing high-affinity bivNb blocking the RBD:ACE2 interaction site demonstrates for the first time a multiplex, antigen-resolved analysis of the presence of neutralizing human IgGs in convalescent individuals suffering from SARS-CoV-2 infection. Compared to other tests analyzing a neutralizing immune response, the NeutrobodyPlex can be performed fully automatable and in high-throughput and can readily be applied for large cohort screening. As it requires only non-living and non-infectious viral material, costs and safety conditions can be substantially decreased^19,29^. Furthermore, the NeutrobodyPlex is highly sensitive as low serum dilutions (tested dilution: 1:400) are sufficient for analysis which significantly reduces patient material compared to standard assays. Considering our findings, it is highly conceivable that the NeutrobodyPlex will open unique possibilities for a detailed classification of the individual immune status with regard to the development of protective antibodies and to monitor the efficiency of desperately needed vaccination campaigns.

## Data availability

The data that support the findings of this study are available from the corresponding authors upon reasonable request.

## Authorship Contributions

N.S.M., T.R.W., M.B. and U.R. designed the study; P.D.K., B.T., T.R.W. performed Nb selection and biochemical characterization; H.S., S.N., A.S. immunized the animal; J.H., D.J., M.B., performed the multiplex binding assay; M.G., A.Z. performed HDX-MS experiments; Mo.S., A.N., J.S.W. and K.S.L. organize and provide patient samples; E.O., G.Z. T.S. designed and performed crystallization studies; N.R., M.S. performed viral infection assays; T.R.W., M.B., J.H., M.G., A.Z., N.R., M.S. and U.R. analyzed data and performed statistical analysis. T.R.W., A.D. and U.R. drafted the manuscript; N.S.M., U.R. supervised the study. All authors critically read the manuscript.

## Acknowledgements

This work was supported by the Initiative and Networking Fund of the Helmholtz Association of German Research Centres (grant number SO-96), the European Union’s Horizon 2020 research and innovation programme under grant agreement No 101003480 – CORESMA.. This work has further received funding from the Ministry of Economic Affairs BadenWürttemberg (FKZ 3-4332.62-NMI/68). We thank Florian Krammer for providing expression constructs for SARS-CoV-2 homotrimeric Spike and RBD.

## Materials and Methods

### Expression constructs

For bacterial expression of Nbs, sequences were cloned into the pHEN6 vector^30^, thereby adding a C-terminal His_6_-Tag for IMAC purification as described previously^20,31^. To generate the hetero-bivalent Nb NM1267, cDNAs were amplified by PCR using forward primer 1230Nfor 5’-GGA CGT CTC AAC TCT CAA GTG CAG CTG GTG GAG TC - 3’ and reverse primer 1230Nrev 5’-CAC CAC CGC CAG ATC CAC CGC CAC CTG ATC CTC CGC CTC CTG AGG ACA CGG TGA CCT GGG CCC - 3’ for NM1230 Nb, and forward primer 1226Cfor 5’-GGT GGA TCT GGC GGT GGT GGA AGT GGT GGC GGA GGT AGT CAG GTG CAG CTG GTG GAA T - 3’ and reverse primer 1226Crev 5’-GGG GAA TTC AGT GAT GGT GAT GGT GGT GTG AGG ACA CGG TGA CCG GGG CC - 3’ for NM1226 Nb. Nbs NM1230 and NM1226 were genetically fused by fusion PCR using forward primer 1230Nfor and reverse primer 1226Crev. Full length cDNA was cloned into Esp3I and EcoRI site of pCDNA3.4 expression vector with N-terminal signal peptide (MGWTLVFLFLLSVTAGVHS) for secretory pathway that comprises Esp3I site.

The pCAGGS plasmids encoding the stabilized homotrimeric spike protein and the receptor binding domain (RBD) of SARS-CoV-2 were kindly provided by F. Krammer^22^. The cDNA encoding the S1 domain (aa 1 - 681) of the SARS-CoV-2 spike protein was obtained by PCR amplification using the forward primer S1_CoV2-for 5’-CTT CTG GCG TGT GAC CGG - 3’ and reverse primer S1_CoV2-rev 5’ - GTT GCG GCC GCT TAG TGG TGG TGG TGG TGG TGG GGG CTG TTT GTC TGT GTC TG - 3’ and the full length SARS-CoV-2 spike cDNA as template and cloned into the XbaI/ NotI-digested backbone of the pCAGGS vector, thereby adding a C-terminal His_6_-Tag. All expression constructs were verified by sequence analysis.

### Nb libraries

Alpaca immunizations with purified RBD and Nb-library construction were carried out as described previously^32^. Animal immunization was approved by the government of Upper Bavaria (Permit number: 55.2-1-54-2532.0-80-14). In brief, nine weeks after immunization of an animal (*Vicugna pacos*) with either C-terminal histidine-tagged RBD (RBD-His_6_), ∼100 ml blood were collected and lymphocytes were isolated by Ficoll gradient centrifugation using the Lymphocyte Separation Medium (PAA Laboratories GmbH). Total RNA was extracted using TRIzol (Life Technologies) and mRNA was reverse transcribed to cDNA using a First-Strand cDNA Synthesis Kit (GE Healthcare). The Nb repertoire was isolated in 3 subsequent PCR reactions using following primer combinations (1) CALL001 (5’-GTC CTG GCT GCT CTT CTA CA A GG-3’) and CALL002 (5’-GGT ACG TGC TGT TGA ACT GTT CC-3’) (2) forward primer set FR1-1, FR1-2, FR1-3, FR1-4 (5’-CAT GGC NSA NGT GCA GCT GGT GGA NTC NGG NGG-3’, 5’-CAT GGC NSA NGT GCA GCT GCA GGA NTC NGG NGG-3’, 5’-CAT GGC NSA NGT GCA GCT GGT GGA NAG YGG NGG-3’, 5’-CAT GGC NSA NGT GCA GCT GCA GGA NAG YGG NGG-3’) and reverse primer CALL002 and (3) forward primer FR1-ext1 and FR1-ext2 (5’-GTA GGC CCA GCC GGC CAT GGC NSA NGT GCA GCT GGT GG-3’, 5‘-GTA GGC CCA GCC GGC CAT GGC NSA NGT GCA GCT GCA GGA-3’ A-) and reverse primer set FR4-1, FR4-2, FR4-3, FR4-4, FR4-5 and FR4-6 (5‘-GAT GCG GCC GCN GAN GAN ACG GTG ACC NGN RYN CC-3’. 5‘-GAT GCG GCC GCN GAN GAN ACG GTG ACC NGN GAN CC-3’. 5‘-GAT GCG GCC GCN GAN GAN ACG GTG ACC NGR CTN CC-3’. 5‘-GAT GCG GCC GCR CTN GAN ACG GTG ACC NGN RYN CC-3’. 5‘-GAT GCG GCC GCR CTN GAN ACG GTG ACC NGN GAN CC-3’. 5‘-GAT GCG GCC GCR CTN GAN ACG GTG ACC NGR CTN CC-3’) introducing SfiI and NotI restriction sites. The Nb library was subcloned into the SfiI/ NotI sites of the pHEN4 phagemid vector^30^.

### Nb Screening

For the selection of RBD-specific Nbs two consecutive phage enrichment rounds were performed. TG1 cells containing the ‘immune’-library in pHEN4 were infected with the M13K07 helper phage, hence the V_H_H domains were presented superficially on phages. For each round, 1 x 10^11^ phages of the ‘immune’-library were applied on RBD either directly coated on immunotubes (10 µg/ml) or biotinylated RBD (5 µg/ml) immobilized on 96-well plates pre-coated with Streptavidin. In each selection round, extensive blocking of antigen and phages was performed by using 5% milk or BSA in PBS-T and with increasing panning round, PBS-T washing stringency was intensified. Bound phages were eluted in 100 mM tri-ethylamin, TEA (pH 10.0), followed by immediate neutralization with 1 M Tris/HCl (pH 7.4). For phage preparation for following rounds, exponentially growing TG1 cells were infected and spread on selection plates. Antigen-specific enrichment for each round was monitored by comparing colony number of antigen vs. no antigen selection. Following panning 492 individual clones of the second selection round were screened by standard Phage-ELISA procedures using a horseradish peroxidase-labeled anti-M13 monoclonal antibody (GE-Healthcare).

### Protein expression and purification

RBD-specific Nbs were expressed and purified as previously described^20,31^. For the expression of SARS-CoV-2 proteins (RBD, stabilized homotrimeric spike and S1 domain), Expi293 cells were used^18^ and the hetero-bivalent Nb NM1267was expressed using the ExpiCHO system. For quality control, all purified proteins were analyzed via SDS-PAGE according to standard procedures. For immunoblotting, proteins were transferred on nitrocellulose membrane (Bio-Rad Laboratories) and detection was performed using anti-His antibody (Penta-His Antibody, #34660, Qiagen) followed by donkey-anti-mouse antibody labeled with AlexaFluor647 (Invitrogen) using a Typhoon Trio scanner (GE-Healthcare, Freiburg, Germany; excitation 633 nm, emission filter settings 670 nm BP 30).

### Biophysical biolayer interferometry (BLI)

To analyze the binding affinity of purified Nbs towards the RBD, biolayer interferometry (BLItz, ForteBio) was performed as per the manufacturer’s protocols. Briefly, biotinylated RBD was immobilized on single-use high-precision streptavidin biosensors (SAX). Depending on the affinity of the RBD-Nb interaction, an appropriate concentration range (15.6 nM - 2 µM) of Nbs was used. For each run, four different Nb concentrations were measured as well as a reference run using PBS instead of Nb in the association step. As negative control, GFP-Nb (500 nM) was applied in the binding studies. Global fits were determined using the BLItzPro software and the global dissociation constant (K_D_) was calculated.

### Bead-based multiplex binding/ competition assay

Purified RBD, S1 domain and homotrimeric spike of SARS-CoV-2 were covalently immobilized on spectrally distinct populations of carboxylated paramagnetic beads (MagPlex Microspheres, Luminex Corporation, Austin, TX) using 1-ethyl-3-(3-dimethylaminopropyl)carbodiimide (EDC)/ sulfo-N-hydroxysuccinimide (sNHS) chemistry. For immobilization, a magnetic particle processor (KingFisher 96, Thermo Scientific, Schwerte, Germany) was used. Bead stocks were vortexed thoroughly and sonicated for 15 seconds. Subsequently, 83 µL of 0.065% (v/v) Triton X-100 and 1 mL of bead stock containing 12.5 x 10^7^ beads of one single bead population were pipetted into each well. The beads were then washed twice with 500 µL of activation buffer (100 mM Na_2_HPO_4_, pH 6.2, 0.005% (v/v) Triton X-100) and activated for 20 min in 300 µL of activation mix containing 5 mg/mL EDC and 5 mg/mL sNHS in activation buffer. Following activation, the beads were washed twice with 500 µL of coupling buffer (500 mM MES, pH 5.0, 0.005% (v/v) Triton X-100) and then proteins were added to the activated beads and incubated for 2 h at 21 °C to immobilize the antigens on the surface. Protein-coupled beads were washed twice with 800 µL of wash buffer (1x PBS, 0.005 % (v/v) Triton X-100) and were finally resuspended in 1,000 µL of storage buffer (1x PBS, 1 % (w/v) BSA, 0.05% (v/v) ProClin). The beads were stored at 4°C until further use. For bead-based multiplex assays, individual bead populations were combined into a bead mix. For the bead-based ACE2 competition binding assay, Nbs were incubated with the bead-mix (containing beads coupled with SARS-CoV-2 homotrimeric spike, RBD and S1 proteins) and biotinylated ACE2 (Sino Biological) which competes for the binding of SARS-CoV-2 spike-derived antigens. Single Nbs or bivNbs were pre-diluted to a concentration of 6.3 µmol/L per Nb in assay buffer. Afterwards, a 4-fold dilution series was made over eight steps in assay buffer containing 160 ng/mL biotinylated ACE2. Subsequently, 25 µL of every dilution was transferred to 25 µL bead-mix in a 96-well half-area plate. The plate was incubated for 2 h at 21 °C, shaking at 750 rpm. Beads were washed using a microplate washer (Biotek 405TS, Biotek Instruments GmbH) to remove unbound ACE2 or Nbs. R-phycoerythrin (PE)-labeled streptavidin was added and incubated for 45 min at 21 °C shaking at 750 rpm to detect biotinylated ACE2 that bound to the immobilized. Afterwards, the beads were washed again to remove unbound PE-labeled streptavidin. Measurements were performed with a FLEXMAP 3D instrument using the xPONENT Software version 4.3 (settings: sample size: 80 µL, 50 events, Gate: 7,500 – 15,000, Reporter Gain: Standard PMT).

### NeutrobodyPlex: Bead-based multiplex neutralizing antibody detection assay

Based on the recently described automatable multiplex immunoassay^23^, the NeutrobodyPlex was developed and similar assay conditions were applied. For the detection of neutralizing serum antibodies, the bead-mix containing beads coupled with purified RBD (receptor-binding domain), S1 (S1 domain), spike (homotrimeric spike), S2 (S2 domain) or N (nucleocapsid) of SARS-CoV-2 was incubated with bivNb NM1257 (concentrations ranging from 1 µM to 6.1 pM) and serum samples of convalescent SARS-CoV-2 patients and healthy donors at a 1:400 dilution. As positive control and maximal signal detection per sample, serum only was included. Bound serum IgGs were detected via anti-human-IgG-PE as previously described ^23^.

### Hydrogen-Deuterium Exchange

#### RBD Deuteration Kinetics and Epitope Elucidation

RBD (5 µL, 73 µM) was either incubated with PBS or RBD-specific Nbs (2.5 µL, 2.5 mg/mL in PBS) at 25 °C for 10 min. Deuterium exchange of the pre-incubated nanobody-antigen complex was initiated by dilution with 67.5 µL PBS (150 mM NaCl, pH 7.4) prepared with D_2_O and incubation for 5 and 50 min respectively at 25 °C. To ensure a minimum of 90% of complex formation, the molar ratio of antigen to Nbs was calculated as previously described^33^, using the affinity constants of 1.37 nM (NM1228), 3.66 nM (NM1226), 3.82 nM (NM1223), 8.23 nM (NM1230) and 8.34 nM (NM1224) (pre-determined by BLI analysis). The final D_2_O concentration was 90%. After 5 and 50 min at 25 °C, aliquots of 15 µL were taken and quenched by adding 15 µL ice-cold quenching solution (0.2 M TCEP with 1.5% formic acid and 4 M guanidine HCl in 100 mM ammonium formate solution pH 2.2) resulting in a final pH of 2.5. Quenched samples were immediately snap-frozen. The immobilized pepsin was prepared by adding 60 µl of 50% slurry (in ammonium formate solution pH 2.5) to a tube and dried by centrifugation at 1000 x g for 3 min at 0 °C and discarding the supernatant. Before injection, aliquots were thawed and added to the dried pepsin beads. Proteolysis was performed for 2 min in a water ice bath followed by filtration using a 22 µm filter and centrifugation at 1000 x g for 30 s at 0 °C. Samples were immediately injected into a LC-MS system. Undeuterated control samples were prepared under the same conditions using H_2_O instead of D_2_O. The same protocol was applied for the Nbs without addition of RBD as well to create a list of peptic peptides. The HDX experiments of the RBD-Nb-complex were performed in triplicates. The back-exchange of the method as estimated using a standard peptide mixture of 14 synthetic peptides was 24%.

#### Chromatography and Mass Spectrometry

HDX samples were analyzed on a LC-MS system comprised of RSLC pumps (UltiMate 3000 RSLCnano, Thermo Fisher Scientific, Dreieich, Germany), a chilling device for chromatography (MéCour Temperature Control, Groveland, MA, USA) and a mass spectrometer Q Exactive (Thermo Fisher Scientific, Dreieich, Germany). The chilling device contained the LC column (ACQUITY BEH C18, 1.7 µm, 300 Å, 1 mm x 50 mm (Waters GmbH, Eschborn, Germany)), a cooling loop for HPLC solvents, a sample loop, and the injection valve and kept all components at 0 °C. Samples were analyzed using a two-step 20 min linear gradient with a flow rate of 50 µL/min. Solvent A was 0.1% (v/v) formic acid and solvent B was 80% acetonitrile (v/v) with 0.1% formic acid (v/v). After 3 min desalting at 10% B, a 9 min linear gradient from 10 to 25% B was applied followed by an 8 min linear gradient from 25 to 68.8%. Experiments were performed using a Q Exactive (Thermo Fisher Scientific, Dreieich, Germany) with 70,000 resolutions instrument configurations as follows: sheath gas flow rate of 25; aux gas flow rate of 5; S-lens RF level of 50, spray voltage of 3.5 kV and a capillary temperature of 300 °C.

#### HDX Data Analysis

A peptic peptide list containing peptide sequence, retention time and charge state was generated in a preliminary LC-MS/MS experiment. The peptides were identified by exact mass and their fragment ion spectrum using protein database searches by Proteome Discoverer v2.1.0.81 (Thermo Fisher Scientific, Dreieich, Germany) and implemented SEQUEST HT search engine. The protein database contained the RBD and the pepsin sequences. Precursor and fragments mass tolerance were set to 6 ppm and 0.05 Da, respectively. No enzyme selectivity was applied, however, identified peptides were manually evaluated to exclude peptides originated through cleavage after arginine, histidine, lysine, proline and the residue after proline ^34^. FDR was estimated using q-values calculated by Perculator and only peptides with high-confidence identification (q-value ≤ 0.01) were included to the list. Peptides with overlapping mass, retention time and charge in Nb and antigen digest, were manually removed. The deuterated samples were recorded in MS mode only and the generated peptide list was imported into HDExaminer v2.5.0 (Sierra Analytics, Modesto, CA, USA). Deuterium uptake was calculated using the increase of the centroid mass of the deuterated peptides. HDX could be followed for 79% of the RBD amino acid sequence. The calculated percentage deuterium uptake of each peptide between RBD-Nb and RBD-only were compared. Any peptide with uptake reduction of 5% or greater upon Nb binding was considered as protected.

### Crystallization and Structural Analysis

#### Production of the RBD domain and complex formation with nanobodies for crystallization

The RBD domain was produced and purified as described previously^18^. As a variation to the established protocol, the production of the RBD (residues 319-541) was performed in the Expi293F™ GntI-expression system. Expi293F™ GntI-cells were cultivated (37 °C, 125 rpm, 8% (v/v) CO_2_) to a density of 5.5×10^6^ cells/mL. The cells were diluted with Expi293F expression medium to a density of 3.0×10^6^ cells/mL, followed by transfection of RBD plasmid (1 µg per mL cell culture) with Expifectamine dissolved in Opti-MEM medium, according the manufacturer’s instructions. 20 h post transfection, the transfection enhancers were added as documented in the Expi293F™ GntI-cells manufacturer’s instructions. The cell suspension was cultivated for 5 days (37 °C, 125 rpm, 8% (v/v) CO_2_) and centrifuged (4 °C, 23900x g, 20 min) to clarify the supernatant. The supernatant was supplemented with His-A buffer stock solution (final concentration in the medium: 20 mM Na_2_HPO_4_, 300 mM NaCl, 20 mM imidazole, pH 7.4), before the solution was applied to a HisTrap FF crude column. The column was extensively washed with His-buffer-A (20 mM Na_2_HPO_4_, 300 mM NaCl, 20 mM imidazole, pH 7.4) before the RBD was eluted with 280 mM imidazole from the column. The RBD was dialyzed against PBS and concentrated to 2 mg/ml. The nanobody complex was formed by mixing the RBD with the purified NM1230 in a molar ratio of 1:1.1, followed by incubation for 3 h at 4°C. For crystallization, the complex was treated with EndoHf to truncate the oligosaccharide chain of the RBD. Therefore, EndoHf (300 U per mg RBD) was added to the RBD:NM1230 complex and incubated for 2 days at 8°C. EndoHf was removed by passing the sample through a MBP-Trap column. Finally, a size-exclusion chromatography using a SD200 16/60 column exchanged the buffer (20 mM HEPES, 150 mM NaCl, pH 7.4) and separated the RBD:NM1230 domain from aggregates and the nanobody excess.

#### Crystallization

The RBD:NM1230 complex was concentrated to 23.2 mg/mL prior to crystallization. Initial crystallization screening was performed on an ART Robbins Gryphon crystallization robot with placing 400 nL drop of RBD:NM1230 and mixed in a 1:1 ratio with the reservoir solution. Initial crystals appeared overnight in crystallization buffer (200 mM MgCl_2_, 20% (w/v) PEG 3350, pH 5.9) and grew to a final size of 30×30×120µm^3^ within 4 days. The crystals were harvested and frozen in liquid nitrogen until data collection.

#### Structure determination and refinement

Data collection was performed on beamline X06SA at the Swiss Light Source (Villigen, Switzerland). For data reduction, the XDS package was used^35^ and the resulting data set was analyzed by XDSSTAT^36^ to check for radiation damage. The tetragonal crystals diffracting to 2.9 Å were initially processed in space group P4_3_2_1_2 containing two copies of RBD:NM1230. The structure was solved by molecular replacement using PHASER^37^ and CHAINSAW^38^ modified templates of the RBD domain (pdb code: 6Z1Z) and a structure homologue of the nanobody (pdb code: 6XC4). Initial refinement involved simulated annealing as implemented in PHENIX^39^ to reduce model bias. Further refinement was done in a cyclic procedure of reciprocal space refinement as implemented in REFMAC5^40^ and real space corrections using COOT^41^. Several cycles of refinement yielded to a model with good stereochemistry and acceptable R-factors of 26.7% and 30.7% for R_work_ and R_free_, respectively (table SY). The structure was validated with MOLPROBITY ^42^ prior deposition to the protein data bank (pdb code 7B27). Figures were generated with PYMOL (SCHRODINGER, L. L. C. The PyMOL molecular graphics system. *Version*, 2010, 1. Jg., Nr. 5, S. 0.) and structure comparison was performed with DALI^43^.

### Cell culture

Caco-2 (Human Colorectal adenocarcinoma) cells were cultured at 37°C with 5% CO_2_ in DMEM containing 10% FCS, 2 mM l-glutamine, 100 μg/ml penicillin-streptomycin and 1% NEAA.

### Viruses

All experiments associated with the SARS-CoV-2 virus were conducted in Biosafety Level 3 laboratory. The recombinant SARS-CoV-2 expressing mNeonGreen (icSARS-CoV-2-mNG) (PMID: 32289263) was obtained from the World Reference Center for Emerging Viruses and Arboviruses (WRCEVA) at the UTMB (University of Texas Medical Branch). To generate icSARS-CoV-2-mNG stocks, 200.000 Caco-2 cells were infected with 50 µl of virus in a 6-well plate, the supernatant was harvested 48 hpi, centrifuged, and stored at -80°C. For MOI determination, a titration using serial dilutions of the mNeonGreen (icSARS-CoV-2-mNG) was conducted. The number of infectious virus particles per ml was calculated as the (MOI × cell number)/ (infection volume), where MOI = −ln(1 − infection rate).

### Viral infection assay

For neutralization experiments, 1 ×10^4^ Caco-2 cells/well were seeded in 96-well plates the day before infection in media containing 5% FCS. Caco-2 cells were co-incubated with the SARS-CoV-2 strain icSARS-CoV-2-mNG at a MOI=1.1 and Nbs or serum samples in serial dilutions in the indicated concentrations. 48 hpi cells were fixed with 2% PFA and stained with Hoechst33342 (1 µg/mL final concentration) for 10 minutes at 37°C. The staining solution was removed and exchanged for PBS. For quantification of infection rates, images were taken with the Cytation3 (Biotek) and Hoechst+ and mNG+ cells were automatically counted by the Gen5 Software (Biotek). Infection rate was determined by dividing the number of infected cells through total cell count per condition. Data were normalized to respective virus-only infection control. Inhibitory concentration 50 (IC_50_) was calculated as the half-maximal inhibitory dose using 4-parameter nonlinear regression (GraphPad Prism).

### Patient samples

112 Serum samples of convalescent SARS-CoV-2-infected and 8 uninfected individuals were analyzed in the course of this study. All samples used were de-identified and pre-existing. Ethical consent was granted from the Ethics Commission of the University of Tuebingen under the votum 179/2020/BO2. Samples were classified as SARS-CoV-2 infected, based upon a self-reported positive SARS-CoV-2 RT-PCR result.

### Analyses and Statistics

Graph preparation and statistical analysis was performed using the GraphPad Prism Software (Version 9.0.0).

## Supplementary Information

### Supplementary Figures

**Supplementary Figure 1:**
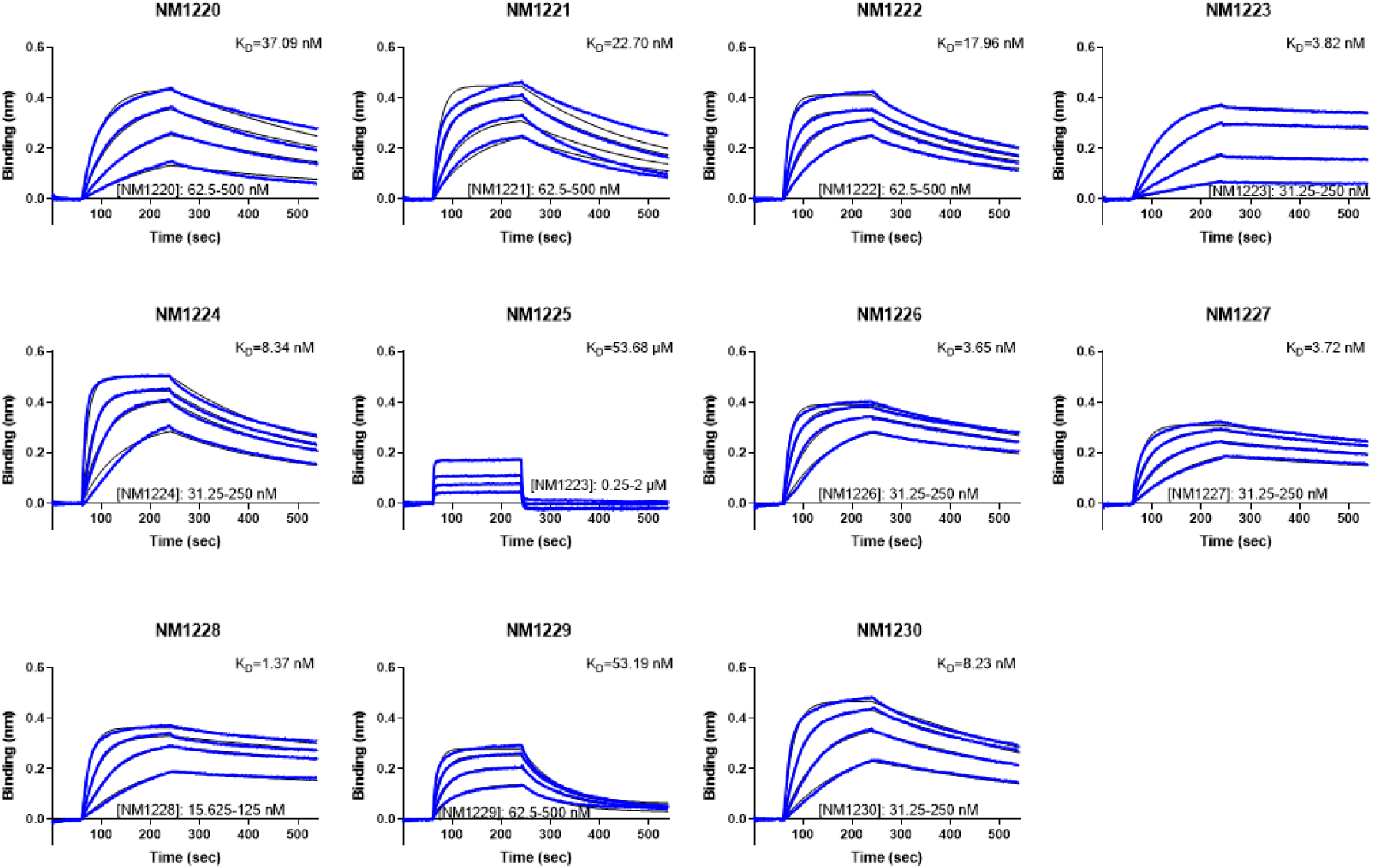
Affinities of RBD-specific Nbs determined by biolayer interferometry. Sensograms of biolayer interferometry-based affinity measurements of 11 identified RBD-specific Nbs are shown. Biotinylated RBD was immobilized on streptavidin biosensors and kinetic measurements were performed by using four concentrations of purified Nbs as indicated.

**Supplementary Figure 2:**
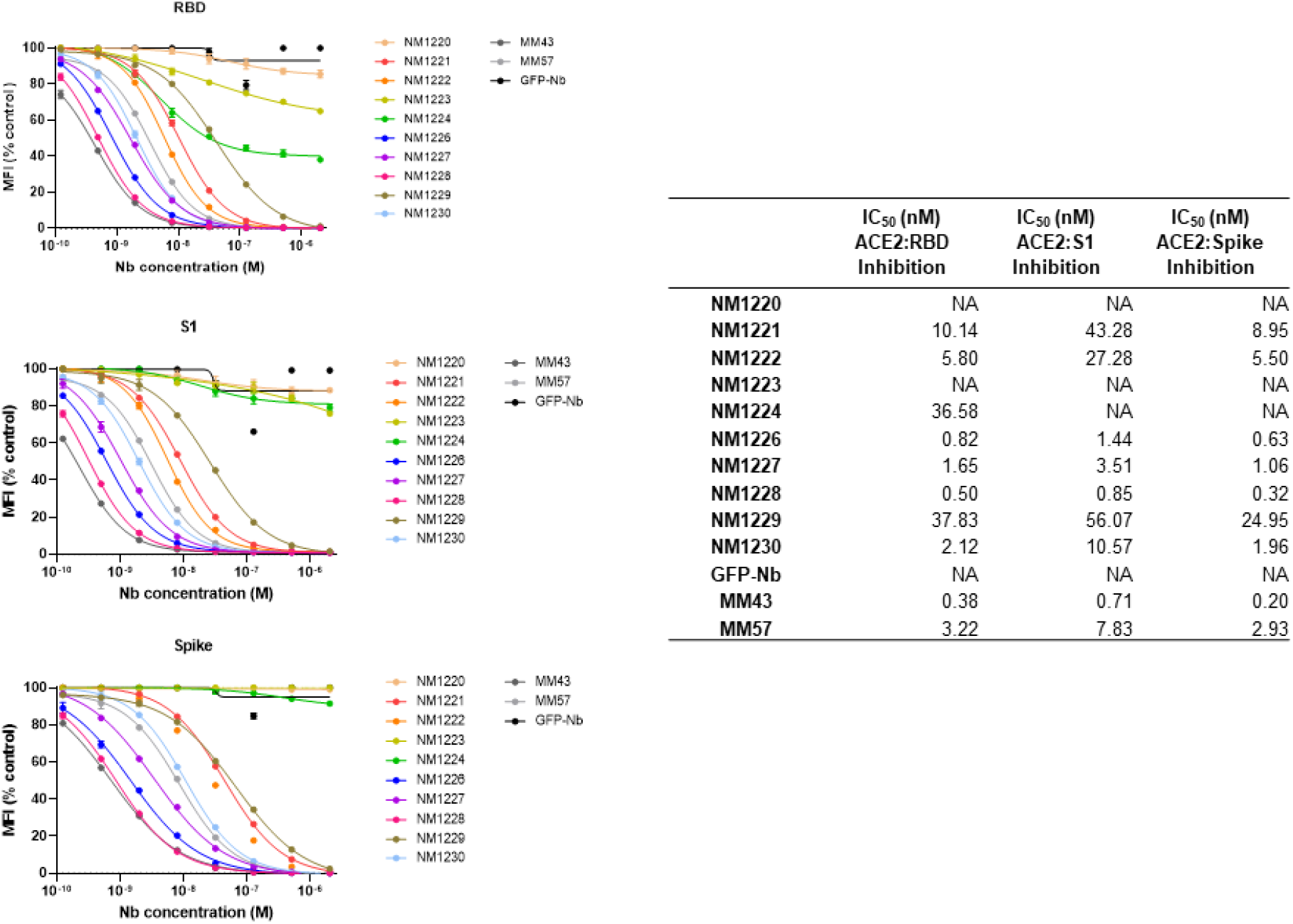
ACE2-competing Nbs identified by multiplex binding assay. Results from multiplex ACE2 competition assay are shown for the three spike-derived antigens: RBD, S1-domain (S1) and homotrimeric spike (Spike). Nbs were diluted from 2.1 µM to 0.12 nM in the presence of 80 ng/mL ACE2 and residual antigen-bound ACE2 was measured. MFI signals were normalized to the maximum detectable signal per antigen given by the ACE2-only control. IC_50_ values were calculated from a four-parametric sigmoidal model. Data are presented as mean +/- SD of three technical replicates (n = 3).

**Supplementary Figure 3:**
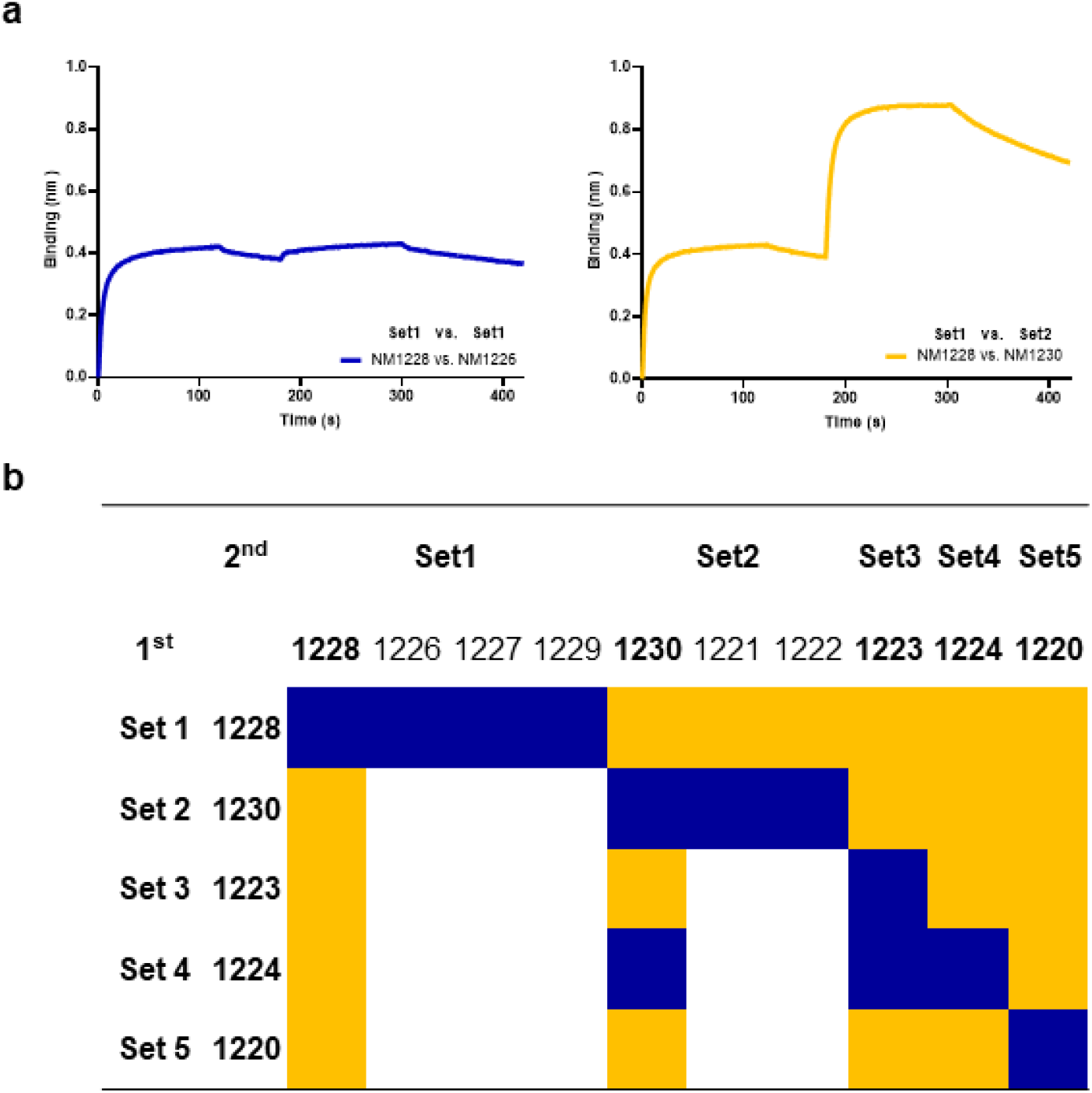
Epitope binning of RBD-specific Nbs. (**a**) Representative sensograms of single biolayer interferometry measurements of Nbs affiliated to the same Nb-Set (NM1228/ NM1226, blue) and two different Nb-Sets (NM1228/ NM1230, orange) are shown. (**b**) Heat map illustration of competitive Nb epitope binning on RBD. Rows and columns represent the loading of the first and second Nb, respectively. Blue colored squares illustrate no additional binding of the second Nb meaning both Nbs belong to the same Nb-Set. Orange colored squares represent additional binding of the second Nb, hence these Nbs belong to different Nb-Sets.

**Supplementary Figure 4:**
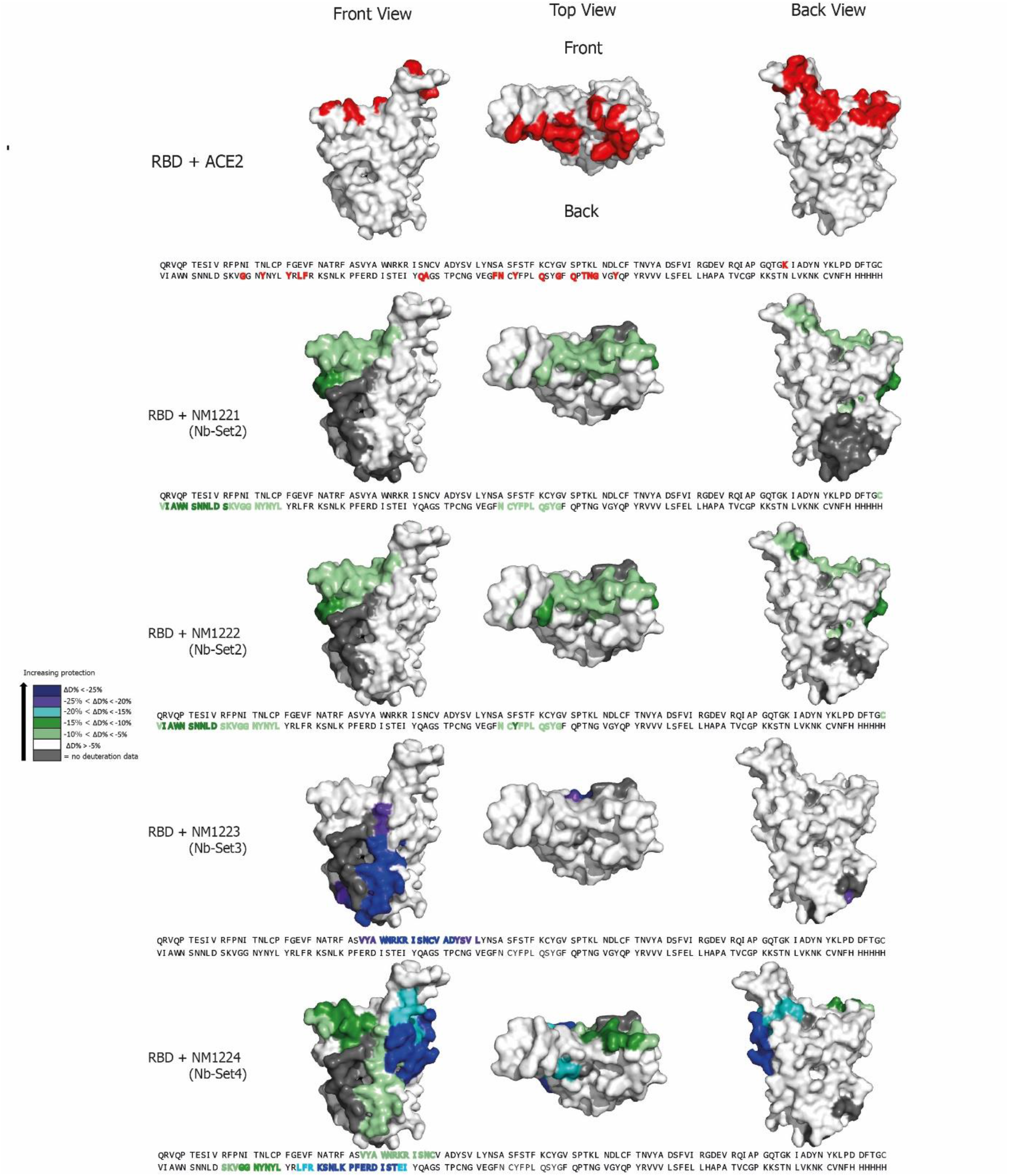
Epitope mapping of Nbs by HDX mass spectrometry. Surface structure model of RBD showing the ACE2 interface and the HDX-MS epitope mapping results of NM1221, NM1222, NM1223 and NM1224. Amino acid residues of RBD (PDB 6M17^2^) involved in the RBD:ACE2 interaction site^2,25^ are shown in red (top panel). RBD epitopes protected upon Nb binding are highlighted in different colors indicating the strength of protection. Accordingly, amino acid residues which are part of the Nb recognized epitopes are highlighted in the RBD sequence.

**Supplementary Figure 5:**
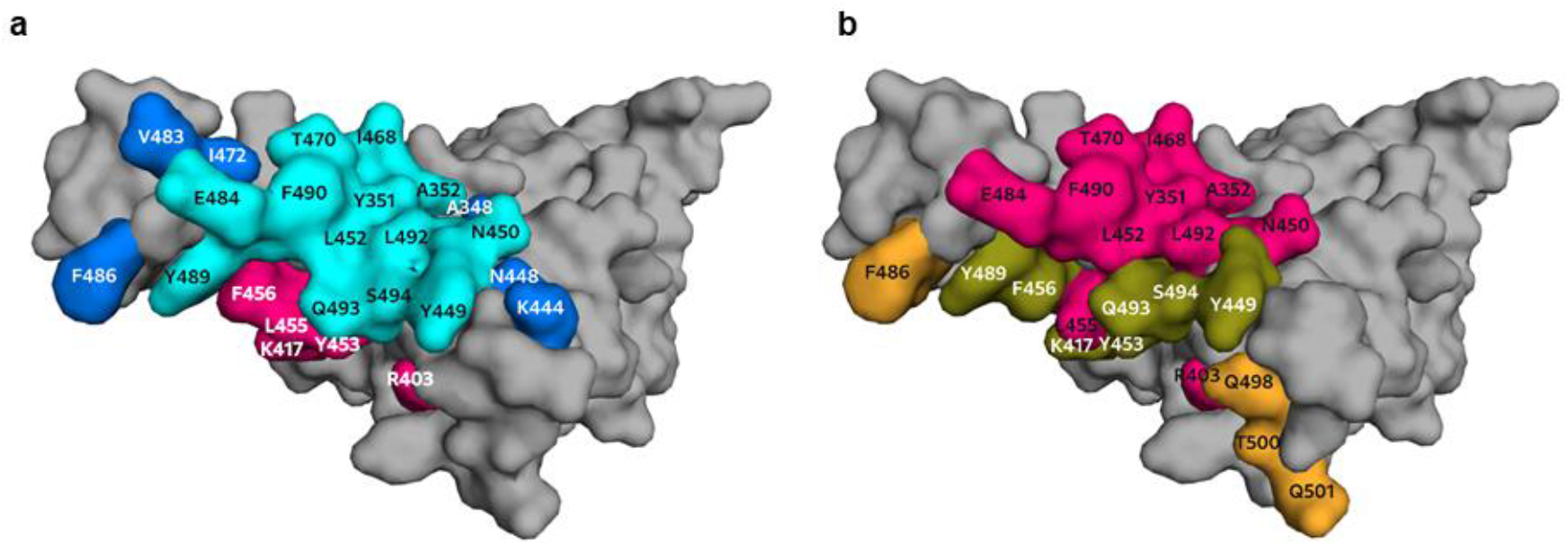
Mapping of RBD-NM1230 interaction sites. (**a**) Individual binding sites of NM1230 (pink) and the Ty1-Nb (blue)^44^ as well as common interaction residues (cyan) are highlighted on the surface representation of RBD. (**b**) Comparison of the ACE2 interaction site and NM1230 epitope on RBD. Common residues are shown in olive, whereas residues exclusively in contact with ACE2 and NM1230 are colored in orange and pink, respectively. All interactions of NM1230, Ty1-Nb and ACE2 are depicted using a distance cut-off of < 4 Å to the surface of RBD.

**Supplementary Figure 6:**
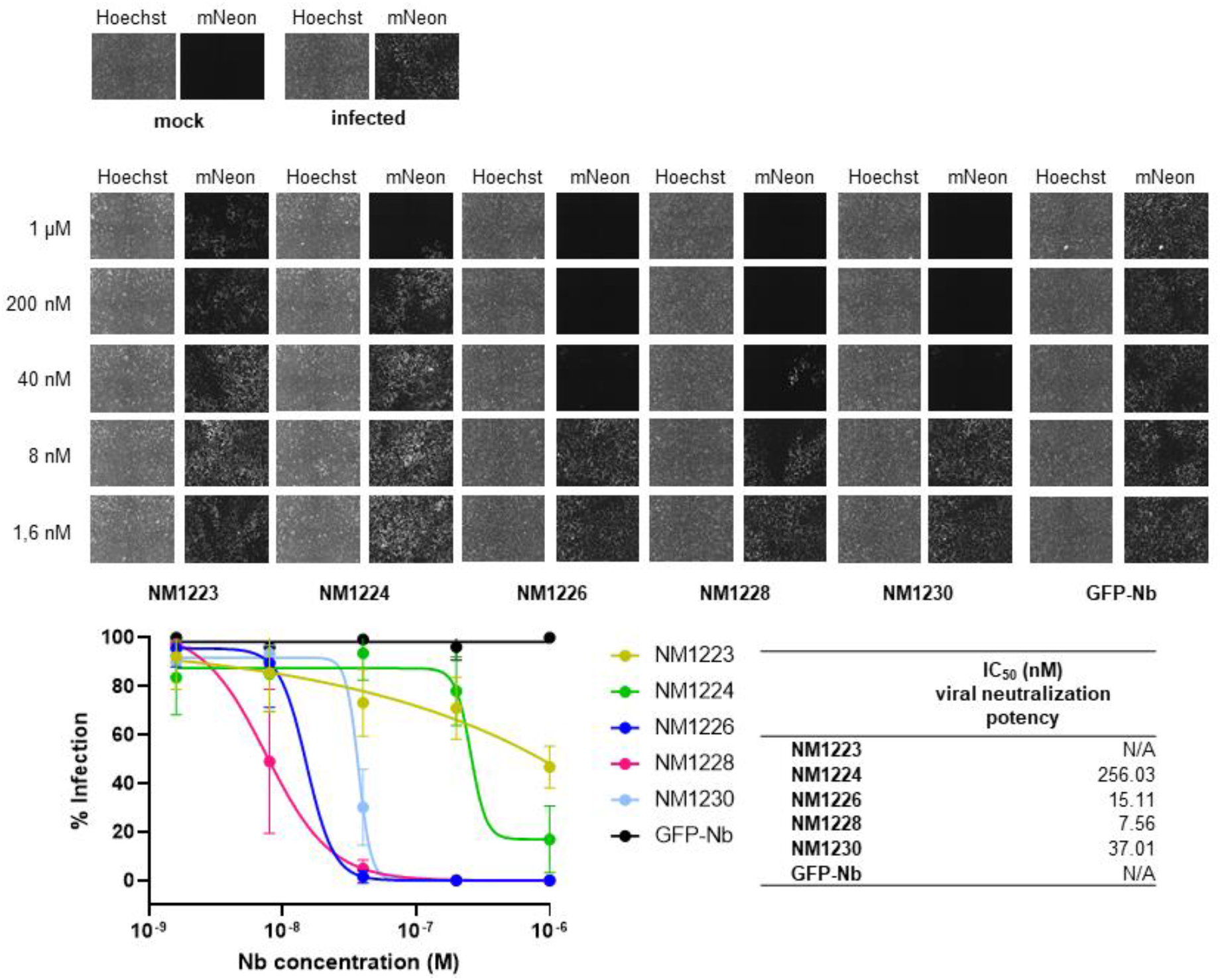
Selected Nbs neutralize SARS-CoV-2 infection. Neutralization potency of NM1223, NM1224, NM1226, NM1228 and NM1230 was analyzed in Caco-2 cells using the SARS-CoV-2-mNG infectious clone. As negative control the GFP-Nb was used. Representative images of human Caco-2 cells upon infection with SARS-CoV-2 expressing mNeonGreen either in presence or absence of serial dilutions of RBD Nbs are shown. Infection rate normalized to virus-only infection control is illustrated as percent of infection (% Infection). IC_50_ value was calculated from a four-parametric sigmoidal model and data are presented as mean +/- SD of three biological replicates (N = 3).

**Supplementary Figure 7:**
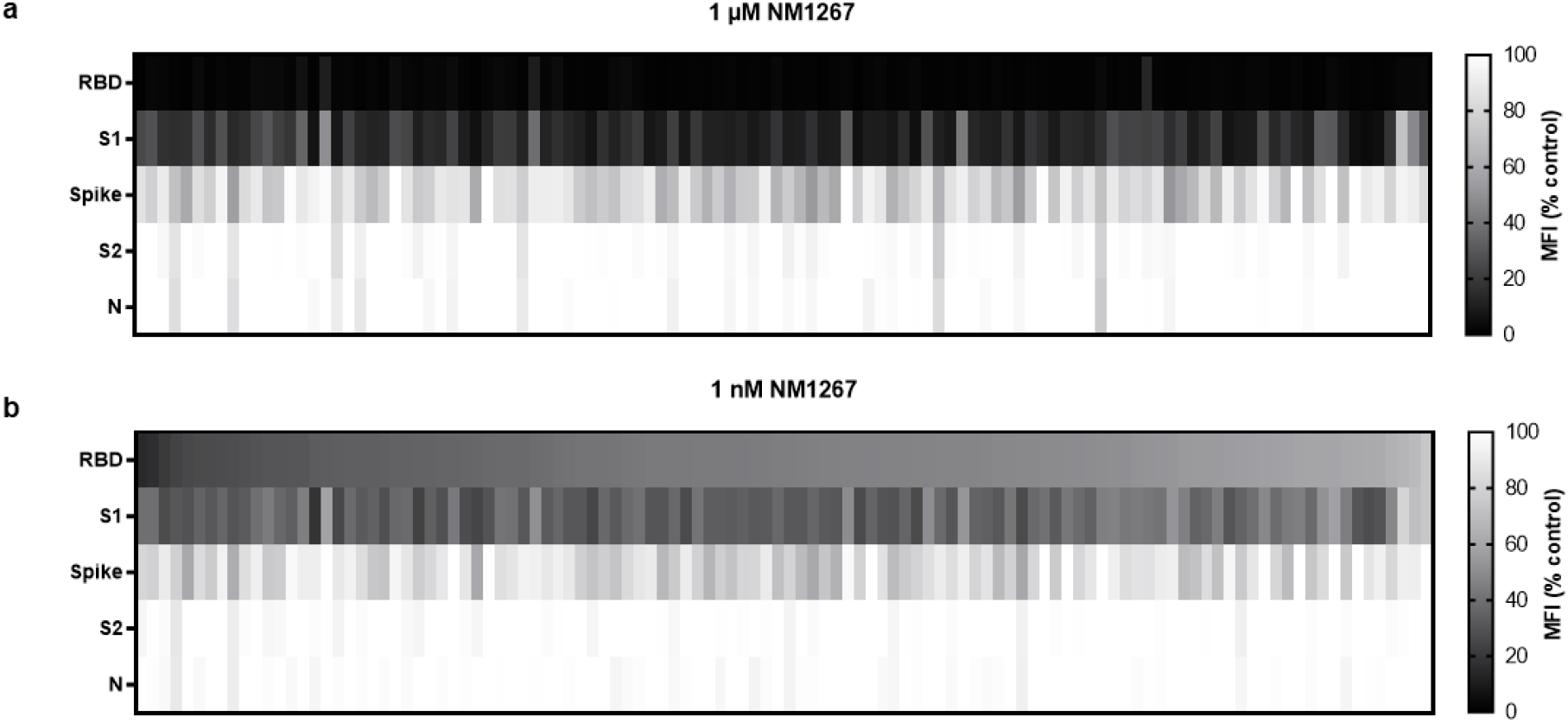
NeutrobodyPlex enables a differentiated analysis on neutralizing IgGs in serum samples of convalescent COVID-19 patients. Serum samples of 112 convalescent SARS-CoV-2-infected individuals were analyzed using the NeutrobodyPlex with antigen-coated beads comprising RBD, S1, spike, S2-domain (S2) and nucleocapsid (N) and two concentrations of NM1267 ((**a**) 1 µM and (**b**) 1 nM). Light-colored squares (high MFI(%control)) are indicative for IgGs outcompeting NM1267 from the RBD:ACE2 interface, dark-colored squares (low MFI (% control) show a continuous displacement of IgGs from serum samples in the presence of NM1267.

**Supplementary Table 1:** **Data collection and refinement statistics for the RBD:NM1230** complex. Values in parentheses are for the highest resolution shell.

|  | **RBD:NM1230 complex** |
| --- | --- |
| **Resolution (Å)** | 46.5-2.90 (2.98-2.90) |
| **Space group** | P4<sub>3</sub>2<sub>1</sub>2 |
| **Unit cell** |  |
| **a, c (Å)** | 63.29, 411.91 |
| **No of reflections** | 112109 (8061) |
| **unique reflections** | 18493 (1340) |
| **Redundancy** | 6.06 |
| **Completeness (%)** | 93.5 (95.7) |
| **I/σ (I)** | 7.8 (1.15) |
| **CC1/2** | 99.8 (66.2) |
| **Wilson B (Å<sup>2</sup>)** | 61 |
| **R<sub>meas</sub> (%)** | 20.7 (186.2) |
| **R<sub>work</sub> / R<sub>free</sub> (%)** | 26.6 / 30.5 |
| **Rmsd** |  |
| **Bond angle (°)** | 1.21 |
| **Bond length (Å)** | 0.016 |
| **Average B-Factor (Å<sup>2</sup>)** |  |
| **RBD (chain A/B)** | 74.6 / 75.2 |
| **NM1230 (chain C/D)** | 64.5 / 81.2 |
| **Ramachandran** |  |
| **Favoured (%)** | 86.3 |
| **Outlier (%)** | 3.9 |

**Supplementary Table 2:** NeutrobodyPlex data illustrated in Figure 6. Value as MFI and MFI (% control).

| # 225 |  |  |  |  |  |  |  |
| --- | --- | --- | --- | --- | --- | --- | --- |
|  | Conc.<br>NM1267<br>[nmol/L] | MFI S1 | MFI S1<br>(% control) | MFI<br>Spike | MFI Spike<br>(% control) | MFI<br>RBD | MFI RBD<br>(% control) |
| NM1267 | 1000.00 | 4453 | 49.58 | 26793 | 69.78 | 103 | 0.49 |
|  | 250.00 | 4425 | 49.27 | 26898 | 70.05 | 162 | 0.77 |
|  | 62.50 | 4377 | 48.74 | 26219 | 68.28 | 336 | 1.60 |
|  | 15.63 | 4423 | 49.24 | 26126 | 68.04 | 1043 | 4.95 |
|  | 3.91 | 4769 | 53.10 | 27975 | 72.85 | 3717 | 17.66 |
|  | 0.98 | 5304 | 59.06 | 28713 | 74.78 | 9823 | 46.66 |
|  | 0.24 | 6556 | 73.00 | 34834 | 90.72 | 16580 | 78.76 |
|  | 0.06 | 8069 | 89.84 | 37128 | 96.69 | 19723 | 93.69 |
| Control n=2<br>(Serum 1:400) | - | 9076 |  | 38978 |  | 21104 |  |
|  | - | 8886 |  | 37819 |  | 20999 |  |
| # 289 |  |  |  |  |  |  |  |
|  | Conc.<br>NM1267<br>[nmol/L] | MFI S1 | MFI S1<br>(% control) | MFI<br>Spike | MFI Spike<br>(% control) | MFI<br>RBD | MFI RBD<br>(% control) |
| NM1267 | 1000.00 | 503.0 | 15.31 | 15943 | 69.07 | 258 | 2.29 |
|  | 250.00 | 512.0 | 15.58 | 16250 | 70.40 | 291 | 2.59 |
|  | 62.50 | 527.5 | 16.05 | 16316 | 70.69 | 349 | 3.10 |
|  | 15.63 | 547.0 | 16.65 | 16319 | 70.70 | 639 | 5.68 |
|  | 3.91 | 641.0 | 19.51 | 16263 | 70.46 | 1667 | 14.82 |
|  | 0.98 | 908.0 | 27.63 | 17369 | 75.25 | 4753 | 42.26 |
|  | 0.24 | 1630.5 | 49.62 | 21303 | 92.29 | 8985 | 79.89 |
|  | 0.06 | 2713.0 | 82.56 | 23808 | 103.15 | 11377 | 100.00 |
| Control n=2<br>(Serum 1:400) | - | 3320.0 |  | 23012 |  | 11250 |  |
|  | - | 3252.5 |  | 23152 |  | 11244 |  |
| # 265 |  |  |  |  |  |  |  |
|  | Conc.<br>NM1267<br>[nmol/L] | MFI S1 | MFI S1<br>(% control) | MFI<br>Spike | MFI Spike<br>(% control) | MFI<br>RBD | MFI RBD<br>(% control) |
| NM1267 | 1000.00 | 1217 | 47.00 | 14574 | 73.63 | 93 | 1.36 |
|  | 250.00 | 1264 | 48.81 | 15273 | 77.16 | 118 | 1.74 |
|  | 62.50 | 1230 | 47.50 | 14896 | 75.26 | 174 | 2.56 |
|  | 15.63 | 1255 | 48.48 | 14788 | 74.71 | 351 | 5.18 |
|  | 3.91 | 1310 | 50.61 | 14993 | 75.75 | 947 | 13.96 |
|  | 0.98 | 1549 | 59.84 | 15985 | 80.76 | 2535 | 37.38 |
|  | 0.24 | 1881 | 72.65 | 17932 | 90.60 | 4648 | 68.52 |
|  | 0.06 | 2225 | 85.96 | 19524 | 98.64 | 6147 | 90.63 |
| Control n=2<br>(Serum 1:400) | - | 2567 |  | 19781 |  | 6666 |  |
|  | - | 2610 |  | 19805 |  | 6899 |  |

| # 272 |  |  |  |  |  |  |  |
| --- | --- | --- | --- | --- | --- | --- | --- |
|  | Conc.<br>NM1267<br>[nmol/L] | MFI S1 | MFI S1<br>(% control) | MFI<br>Spike | MFI Spike<br>(% control) | MFI<br>RBD | MFI RBD<br>(% control) |
| NM1267 | 1000.00 | 159 | 25.37 | 15289 | 86.71 | 48 | 2.05 |
|  | 250.00 | 163 | 25.93 | 14686 | 83.29 | 53 | 2.24 |
|  | 62.50 | 153 | 24.41 | 14857 | 84.26 | 65 | 2.77 |
|  | 15.63 | 163 | 26.01 | 14923 | 84.63 | 116 | 4.95 |
|  | 3.91 | 192 | 30.63 | 14895 | 84.47 | 321 | 13.70 |
|  | 0.98 | 237 | 37.73 | 15036 | 85.27 | 853 | 36.40 |
|  | 0.24 | 335 | 53.45 | 16833 | 95.47 | 1647 | 70.28 |
|  | 0.06 | 531 | 84.72 | 17305 | 98.14 | 2165 | 92.38 |
| Control n=2<br>(Serum 1:400) | - | 611 |  | 17575 |  | 2329 |  |
|  | - | 643 |  | 17690 |  | 2359 |  |
| # 159 |  |  |  |  |  |  |  |
|  | Conc.<br>NM1267<br>[nmol/L] | MFI S1 | MFI S1<br>(% control) | MFI<br>Spike | MFI Spike<br>(% control) | MFI<br>RBD | MFI RBD<br>(% control) |
| NM1267 | 1000.00 | 96 | 49.61 | 12792 | 99.38 | 37 | 5.07 |
|  | 250.00 | 87 | 45.19 | 11708 | 90.96 | 37 | 5.00 |
|  | 62.50 | 90 | 46.75 | 11503 | 89.36 | 43 | 5.83 |
|  | 15.63 | 79 | 41.04 | 11340 | 88.10 | 51 | 6.99 |
|  | 3.91 | 82 | 42.60 | 11263 | 87.50 | 92 | 12.54 |
|  | 0.98 | 86 | 44.68 | 11437 | 88.85 | 257 | 35.23 |
|  | 0.24 | 117 | 60.78 | 12483 | 96.98 | 531 | 72.79 |
|  | 0.06 | 166 | 85.97 | 13044 | 100.00 | 681 | 93.28 |
| Control n=2<br>(Serum 1:400) | - | 191 |  | 13077 |  | 758 |  |
|  | - | 194 |  | 12668 |  | 702 |  |

**Supplementary Table 3:**
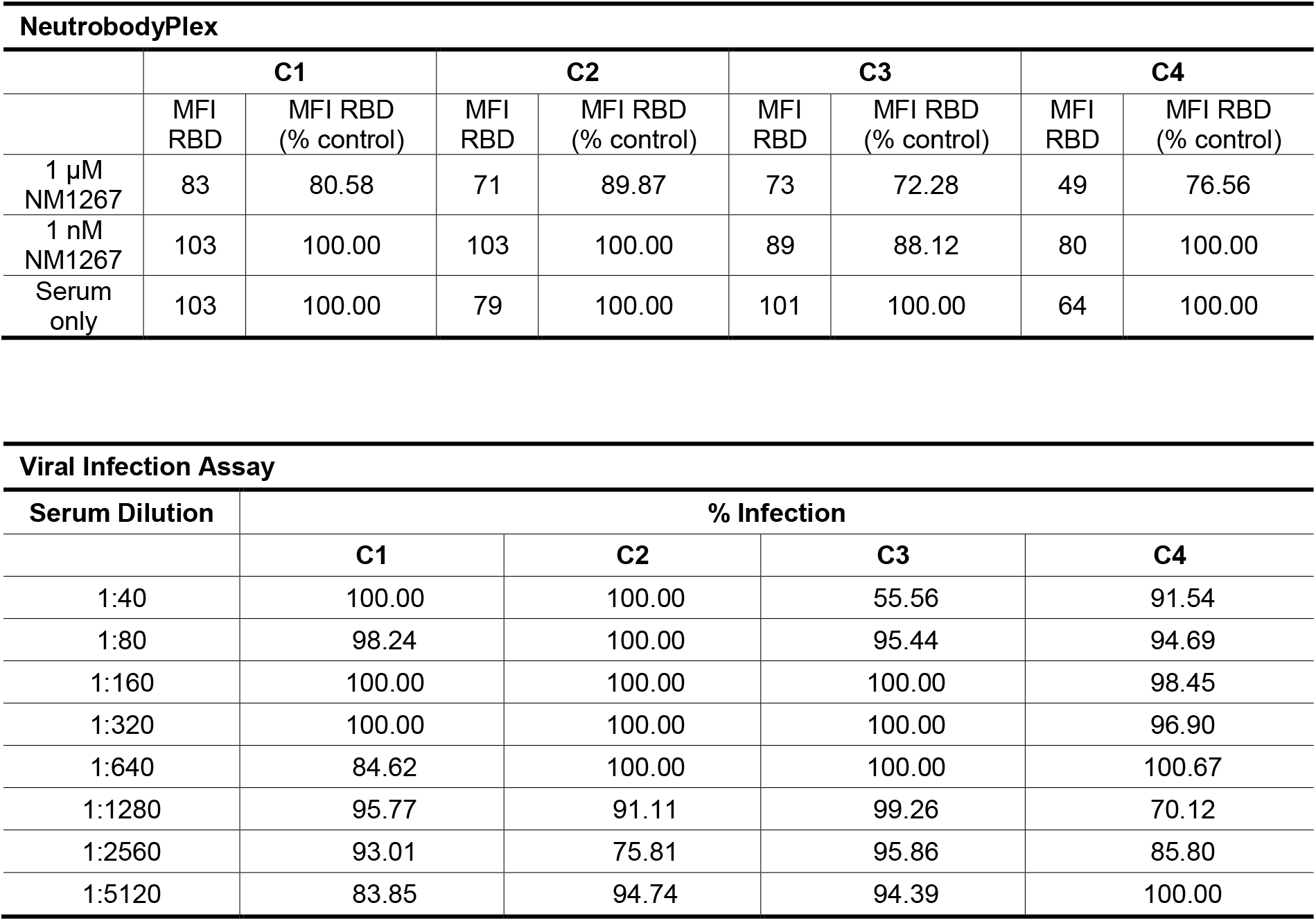
Data of control samples (C1-C4) from healthy donors tested in the NeutrobodyPlex and viral infection assay.

## Notes

### Competing Interest Statement

The authors have declared no competing interest.

### Summary of Updates

In the substantially revised version we added novel data including the crystal structure of Nanobody NM1230 in complex with RBD in the results section. We additionally generated and described a novel bivalent Nanobody (NM1267) as the most potent IgG surrogate which was applied in the described NeutrobodyPlex assay. Finally the NeutrobodyPlex was used to analysed a large patient cohort of 120 individuals (all result section) With Elena Ostertag, Georg Zocher and Thilo Stehle we added three additional authors which performed the RBD:NM1230 crystallization analysis. In the revised version Supplementary Information are added to the main text. All parts were adapted for clarification

